# The dengue virus non-structural protein 1 (NS1) is secreted from mosquito cells in association with the intracellular cholesterol transporter chaperone caveolin complex

**DOI:** 10.1101/370932

**Authors:** Romel Rosales Ramirez, Juan E. Ludert

**Author notes:** Corresponding author: (JEL).

## Abstract

Dengue virus (DENV) is a mosquito-borne virus of the family *Flaviviridae.* The RNA viral genome encodes for a polyprotein that is co-translationally processed into three structural proteins and seven non-structural proteins. The non-structural protein 1 (NS1) is a multifunctional viral protein actively secreted in vertebrate and mosquito cells during DENV infection. In mosquito cells, NS1 is secreted in a caveolin-1 (CAV-1) dependent manner by an unconventional pathway. The caveolin chaperone complex (CCC) is a cytoplasmic complex formed by caveolin-1 and the chaperones FKBP52, Cy40 and CyA which is responsible for cholesterol traffic inside the cell. In this work, we demonstrate that in infected mosquito cells, DENV NS1 is secreted by an early and unconventional route that bypasses the Golgi apparatus in close association with the CCC. Treatment of mosquito cells with classic secretion inhibitors such as brefeldin A, golgicide A and Fli-06 showed no effect on NS1 secretion, but significant reductions in recombinant luciferase secretion and virion release. Silencing the expression of CAV1, FKBP52 with siRNAs or the inhibition of CyA by cyclosporine A resulted in significant decrease in NS1 secretion without affecting virion release. Using co-localization, co-inmunoprecipitation and proximity ligation assays, NS1 was found to co-localize and interact with all the protein components of the CCC in mosquito infected cells. In addition, CAV-1 and FKBP52 expression was found augmented in DENV infected cells. Finally, the treatment of ZIKV infected mosquito cells with brefeldin A and golgicide A showed no effect on NS1 secretion, while affecting virion release. ZIKV NS1 was also found to co-localize with CAV-1 in infected mosquito cells. These results suggest that in mosquito cells, ZIKV NS1 follows the same secretory pathway observed for DENV NS1. The association of NS1 with the cholesterol transporter CCC agrees with the lipoprotein nature of secreted hexameric NS1.

**AUTHOR SUMMARY:** Dengue protein NS1 is secreted in infected mosquito and vertebrate cells. In humans, secreted NS1 have been associated with pathogenesis. In mosquito cells, NS1 follows an unconventional secretion pathway that is dependent on Caveolin-1. This work shows that in mosquito cells, NS1 secretion is associated to the chaperone caveolin complex, a complex formed by caveolin-1 and several chaperones, in charge of cholesterol transport within the cells. Reduction of the expression or the activity of chaperone caveolin complex in mosquito infected cells, diminished the secretion of NS1 without affecting virion release. Direct interaction between NS1 and the chaperone caveolin complex proteins was demonstrated by several assays. Moreover, increased expression of the caveolin-1 and co-chaperone FKBP52 during dengue infection was found, presumably in response to the higher requirements of these proteins during dengue virus infection. Results obtained with ZIKV infected mosquito cells suggest that also ZIKV NS1 is released following an unconventional secretory route in association with the chaperone caveolin complex. The functions of secreted NS1 within mosquito are unclear. However, giving the importance of the soluble NS1 in the vertebrate host, manipulation of the NS1 secretory route may prove a valuable strategy for dengue mosquito control and patient treatment.

## INTRODUCTION

Dengue is an arthropod-borne viral disease caused by any of the four dengue virus serotypes (DENV 1–4), which are transmitted by *Aedes* mosquitoes [1]. Dengue infection may develop into a life-threatening illness. There are currently no specific therapeutics and substantial vector control efforts have not stopped its rapid emergence and global spread. The risk of dengue virus infection and its public health burden are hard to estimate, but at least 100 countries are currently endemic for dengue [2]. DENV has a single stranded RNA genome of positive polarity. The genome of DENV is translated into a single polyprotein and encodes for three structural (C, E, prM/M) and seven non-structural (NS) (NS1, NS2A, NS2B, NS3, NS4A, NS4B, NS5) proteins. The polyprotein is then cleaved by host and viral proteases to release individual viral proteins [3]. The viral genome replication process within the host cell is mainly driven by the NS proteins. The NS1 protein act as an scaffolding protein that anchors the replication complex to the ER membrane and interacts physically with NS4B [4]. The NS1 protein is a 352-amino-acid polypeptide with a molecular weight of 46–55 kDa, depending on its glycosylation status. The NS1 protein exists in multiple oligomeric forms and is found in different cellular locations: a cell membrane-bound form in association with virus-induced intracellular vesicular compartments, on the cell surface and as a soluble secreted hexameric lipoparticle [4]. The NS1 monomeric form rapidly dimerizes in the endoplasmic reticulum (ER), then three dimeric forms of NS1 arrange to form an hexamer [5]. The hexameric form of NS1 shows an open barrel form filled with lipids and cholesterol which resemble the lipid composition of the HDL particle [6].

Recent studies have shown that the DENV NS1 protein was not only secreted from vertebrate cells, but that was also efficiently secreted from mosquito cells lines [7,8]. The secretion of NS1 in vertebrate cells follows the classical Golgi-pathway [9]. However, NS1 secretion in infected mosquito cells is associated to a caveolin-1 (CAV-1) dependent pathway and was found to be Brefeldin A (BFA)-insensitive, suggesting a traffic route that bypasses the Golgi-complex [10].

Caveolae are made up of interlocking heteropolymers of a family of small proteins (caveolins: 1-3) and a second family of accessory structural proteins (flotillins and three family of cavins). The caveolar architecture is connected with unstructured cavin filaments by coiled-coil domains into a polygonal net-like complex. This complex is believed to provide scaffolding for compartmented cellular processes and participates in multiple cellular functions, including endocytosis, transcytosis, membrane homeostasis, inflammation, and signal transduction [11]. Caveolin 1 (Cav-1), a 21–24-kDa scaffolding protein, is not only a key structural component of the caveolae organelle but also plays an important in the transport of free-cholesterol inside the cell [12,13]. The chaperone caveolin complex (CCC) is a cytosolic complex reported to transport cholesterol synthetized *de novo* from the ER to cell membranes or other compartments within the cell. CCC has been described as a complex of CAV-1, Cyclophilin A (CyA), FK506-binding protein 4 or heat shock protein 56 (FKBP52), and Cyclophilin 40 or D (Cy40) [12,14]. Cyclophilin A (CyA), an 18 kDa peptidylprolyl cis-trans isomerase, is a ubiquitous and multifunctional protein. In addition to its role as a host-cell receptor for cyclosporine A, CyA has diverse functions in inflammatory conditions and diseases [15,16]. The 52-kDa FK506-binding protein (FKBP52) an immunophilin belonging to the FKBP family, is a known co-chaperone of the heat shock protein 90 (HSP90) and thus may play a role in the intracellular trafficking of hetero-oligomeric forms of the steroid hormone receptors [17,18]. Cyclophilin 40 (Cy40), a member of a family of highly homologous peptidylprolyl cis-trans isomerases (PPIases), is known to play a role in mitochondrial permeability transition (MPT), being an integral constituent of the MPT pore [19].

Given the CAV-1 dependent secretion of NS1 protein in mosquito cells and the lipoprotein nature of the released hexameric form of NS1, it was found plausible to study the association of NS1 traffic to the *novo* cholesterol transport in DENV infected mosquito cells. In this work, data is presented indicating that in infected mosquito cells, DENV NS1 enters the unconventional secretory pathway very early after maturation in the ER and usurps the cholesterol transport between the ER and the plasma membrane, mediated by the CCC, to reach the extracellular space. In addition, data is presented suggesting that a similar pathway is used for the secretion of Zika Virus NS1 protein in infected mosquito cells.

## RESULTS

### NS1 secretion is not affected by drugs that disrupt early steps of the classical secretion pathway

Golgicide A (GCA) is a powerful inhibitor of the COPI vehicle transport from ER to Golgi [20]. Thus, the cytotoxicity of GCA in the mosquito cell lines (C6/36, Aag2) and the vertebrate cell line BHK-21, used for comparisons, was measured using the reduction of tetrazolium salts to examine proliferation in cells treated with serial dilutions of GCA. No significant cytotoxicity was observed under 30 μM of GCA in any of the three cell types (Fig 1A). Fli-06 is a novel drug which inhibits the diffusion of ER synthetized proteins to the ER-exit sites [21,22]. Fli-06 cytotoxicity was determined in C6/36 and Aag2 cells also using also tetrazolium salt reduction. No significant cytotoxicity was observed under 100 μM of Fli-06 in the mosquito cells lines (Fig 1B). Thus, concentration of 27 μM and 100 μM were used for GCA and Fli-06, respectively, since these concentrations proved non-toxic, yet effective in causing Golgi disruption (Fig 1C). Figures 1D and 1E show that the GCA-treatment of DENV infected C6/36 and Aag2 cells did not cause any change in NS1 secretion. In contrast, NS1 secretion was reduced over 70% in infected BHK-21 cell treated either with GCA or BFA (Fig 1F). As expected, virion release was reduced in mosquito and vertebrate cells after GCA or BFA treatment (Fig 1G, 1H and 1I). To further explore the traffic route of NS1 in DENV infected mosquito cells, cells were also treated with Fli-06. Treatment of infected C6/36 or Aag2 cells with non-toxic concentrations of Fli-06 shows no effect in NS1 secretion, while significantly affecting virion release (Fig 1G, 1H). Experiments with Fli-06 were not carried out in BHK-21 due to high drug toxicity. The results with GCA and Fli-06 indicate NS1 is not secreted by a classical secretory pathway and that newly synthetized DENV NS1 do not reach the ER-exit sites and leaves the ER compartment very early after synthesis.

**Fig 1.**
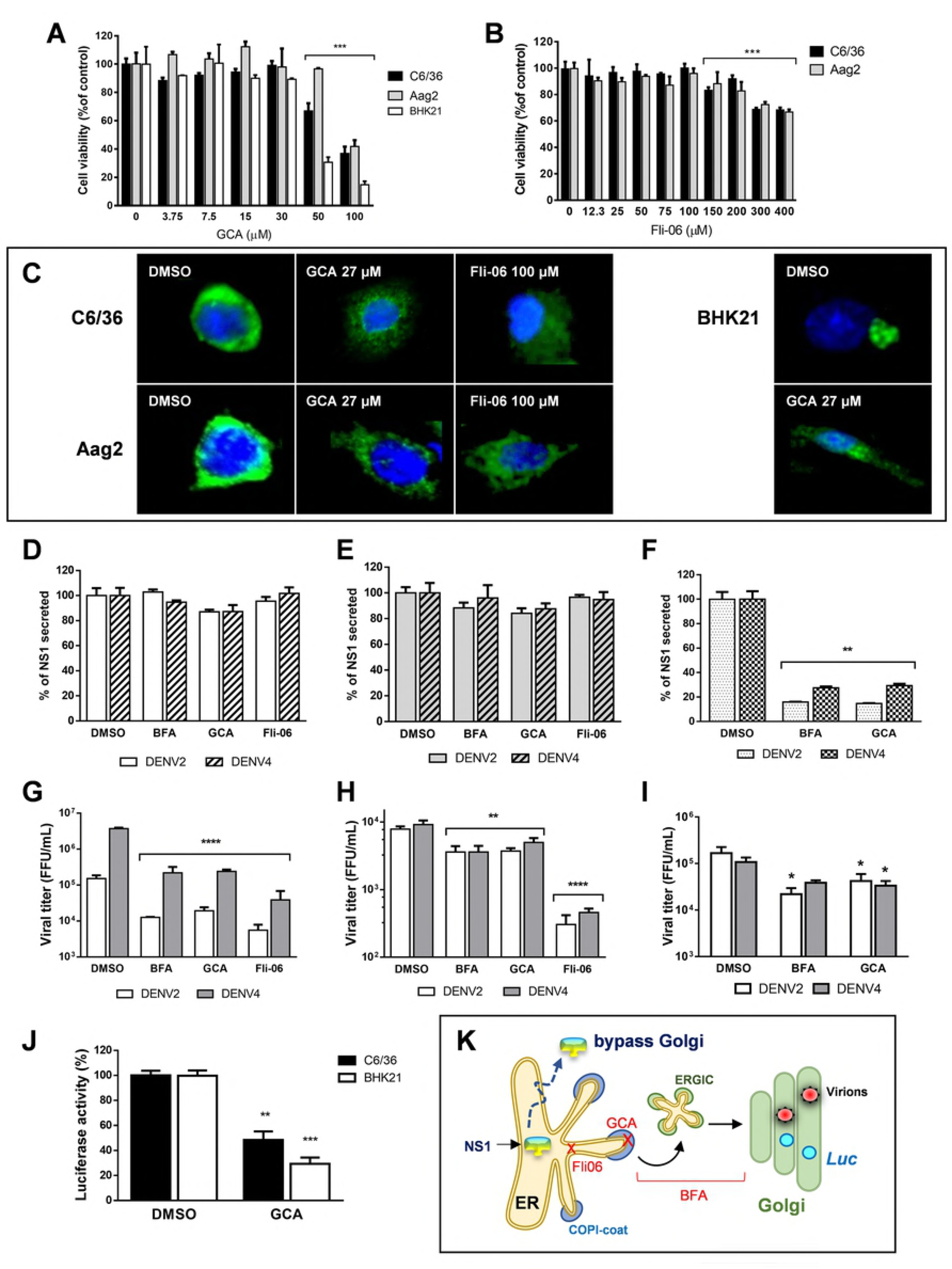
NS1 secretion is not affected in mosquito cells treated with classical secretion pathway inhibitors. **A** Cell toxicity of Golgicide A (GCA) in C6/36, Aag2 and BHK-21 cells. **B** Cell toxicity of Fli-06 in C6/36 and Aag2 cells. ***A*** *and* ***B*** Cells were seeded in 96-well plates and then treated with serial dilutions of GCA and Fli-06 for 24 hours. Cell viability was measure with MTT assay kit and expressed as percentage of control (DMSO-treated). Data are mean of 3 independent experiments ± standard error; significant differences compared with controls are denoted by *(P < 0.0001). **C** Golgi matrix integrity after cell treatment with GCA 27 μM and Fli-06 100 μM for 24 h was determined by staining the Golgi matrix. Cells were fixed and stained with anti-GM130 (cis-Golgi, in green) and DAPI (nuclei in blue). **D** Inhibition of classical secretory pathway in C6/36 cells. **E** Inhibition of classical secretory pathway in Aag2 cells. **F** Inhibition of classical secretory pathway in BHK-21 cells. **G** Viral release in C6/36 cells drug treated. **H** Viral release in Aag2 cells drug treated. **I** Viral release in BHK-21 cells drug treated. ***D, E*** *and* ***F***; Cells were infected with DENV2 or DENV4 at MOI of 3 for 1 h. The cells were washed 3 times and after treated with DMSO (control), 7 μM BFA, 27 μM GCA or 100 μM Fli-06 and supernatants harvested at 24 hpi DENV NS1 was measured using Platelia NS1 Ag (Bio-Rad) and represented as percentage of NS1 secreted compared to DMSO. ***G, H*** *and* ***I***; Viral titers were measured in supernatants from treated cells by focus forming assay in BHK-21 cells following standard protocol. **J** C6/36 cells (pAc5-mCherry-GLuc-neo-transfected) and BHK-21 cells (ptk-GLuc-transfected) expressing *Gaussia* Luciferase were treated with DMSO and GCA 27 μM for 24 h. Luciferase activity was measured in supernatants and represented as percentage of Luc activity. ***D, E, F, G, H, I*** *and* ***J*** The experiments were performed in triplicate, and the bars represent means ± standard error (error bars). Data was evaluated using the 2way ANOVA test and significant differences compared with DMSO treatment group, respectively, are denoted by *(P ≤0.05). **K** Diagram showing the mode of actions of the drugs used and the early ER exit of NS1 in mosquito cells.

Since the release of mature virions, which are multimolecular complexes, may not be equivalent to the release of a single protein, an additional control for protein trafficking through the classical secretory pathway was included. To this aim, a vector, pAc5-mCherry-GLuc-neo, constitutively expressing *Gaussia* Luciferase in C6/36 cell was constructed (S1 Fig). To evaluate the effect of GCA treatment on GLuc secretion, cells were treated with GCA for 24 h and the activity of GLuc present in the cell supernatants analyzed using a luminescence assay. A decrease in luciferase activity, in relation to the control (DMSO), was detected upon treatment of C6/36 with GCA, and in BHK-21 cells transfected with ptk-GLuc vector, run in parallel as control (Fig.1J). These results indicated that the luciferase reporter is secreted in both mosquito and vertebrate cells following a classical secretory route and reinforce the results indicating that in mosquito cells DENV NS1 is secreted by an unconventional secretory route that bypasses the Golgi (Fig 1K).

### NS1 secretion is dependent on the CCC

The CCC is formed by the association of CAV-1 with the chaperones CyA, FKBP52, Cy40 [12,13]. The CCC is responsible for cholesterol transport inside the cell. Previous data by us [10] indicated that NS1 secretion was dependent on CAV-1. Thus, to fully evaluate the participation of the CCC in DENV NS1 secretion from mosquito cells, CyA was pharmacologically inhibited by treatment with Cyclosporin A (CsA) and the expression of the chaperones FKBP52 and Cy40 knocked down using siRNAs. Previously, the non-cytotoxic concentration for CsA in mosquito and vertebrate cells were determined (Fig 2A). DENV2 or DENV4-infected C6/36 or Aag2 cells were treated with 9 μM CsA for 24 hours; NS1 secretion was reduced in about 30% and 40% in C6/36 and Aag2 cells, respectively (Fig 2B, 2E). However, virion release in the treated mosquito cells was not significantly affected (Fig 2C and 2F). In contrast, the inhibition of CyA in BHK-21 cells did not have any effect in either NS1 or virion secretion (Figs 2H and 2I). To further corroborate the participation of CyA in cholesterol and lipid traffic, the lipid droplet count in CsA treated versus untreated cells was determined. In all cases, C6/36, Aag2 and BHK-21 cells treated with CsA showed a diminished amount of lipid droplets per cell as compared with un-treated cells (Fig 2D, 2G and 2J).

**Fig 2.**
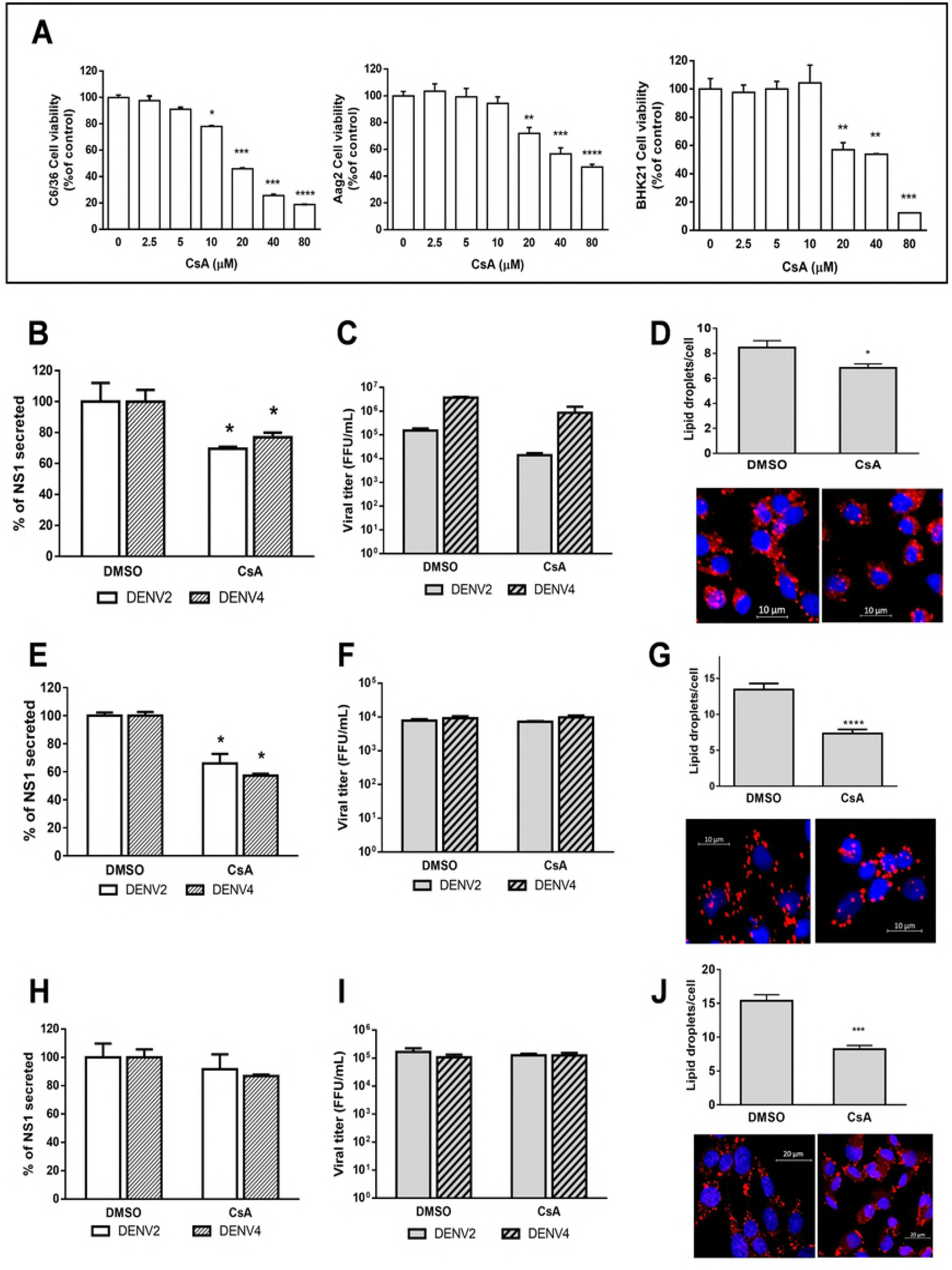
NS1 secretion is inhibited in mosquito cells treated with CsA. **A** Cell toxicity of cyclosporine A (CsA) in C6/36, Aag2 and BHK-21 cells. Cells were seeded in 96-well plates and then treated with serial dilutions of CsA for 24 hours. Cell viability was measure with MTT assay kit and expressed as percentage of control (DMSO-treated). Data are mean of 3 independent experiments ± standard error, and significant differences compared with control are denoted by *(P < 0.0001). **B** NS1 secretion in CsA treated (9 μM) C6/36 cells **C** viral release in CsA treated (9 μM) C6/36 cells. **D** Lipid droplet counting in C6/36 cells treated with CsA. **E** NS1 secretion in CsA treated Aag2 (9 μM) cells. **F** viral release in CsA treated (9 μM) Aag2 cells. **G** Lipid droplet counting in Aag2 cells treated with CsA. **H** NS1 secretion in CsA treated (9 μM) BHK-21 cells **I** viral release in CsA treated (9 μM) BHK-21 cells. **J** Lipid droplet counting in BHK-21 cells treated with CsA. ***B***, ***E*** *and* ***H*** Cells were infected with DENV2 or DEN4 at MOI of 3 for 1 h. The cells were washed 3 times and treated with DMSO (control) or 9μM CsA and harvested at 24 hpi DENV NS1 in supernatants was measured using Platelia NS1 Ag (Bio-Rad) and represented as % of NS1 secreted compared to control (DMSO). ***C***, ***F*** *and* ***I*** Viral titers in supernatants were measured by focus forming assay in BHK-21 cells according to standard procedure. ***D***, ***G*** *and* ***J*** Lipid droplets were stained with Nile red (red) and nuclei with DAPI (blue). LD were counted in maximum projection images in treated cells from at least 25 cells by group (DMSO or CsA) using spot detector plugin using Icy software. LD median count differences were compared using Student’s t test denoted by * (P ≤ 0.05). ***B***, ***C***, ***E***, ***F***, ***H*** *and* ***I*** The experiments were performed in triplicate, and the bars represent means ± standard error (error bars). Data was evaluated using the 2way ANOVA test and significant differences compared with DMSO treatment group, respectively, are denoted by *(P ≤0.05).

The expression of FKBP52 in C6/36 cells was knock down by siRNA transfection. Optimal gene knock down was achieved after 48h post transfection (Fig 3A). The effect of FKPB52 reduced expression in cholesterol and lipid metabolism was evaluated measuring the lipid droplet counting per cell and a reduction of about 50% in the number of lipid droplets was observed after the treatment (Fig 3B). In cells treated with siRNA FKBP52, a significant reduction in NS1 of about 20% secretion was observed (Fig 3C). Viral release was not affected by FKBP52-knock down (Fig 3D). Given that FKBP52 is a co-chaperone of HSP90 [17,23], we wanted to discard if the observed inhibitory effect on NS1 secretion might be related to a reduction in HSP90 activity. Thus, the effect of the HSP90 inhibitor geldanamycin (GA) in NS1 release from DENV infected C6/36 cells was tested. As shown in Figure 3E, NS1 secretion was not affected after GA treatment. The expected reduction in virus yield observed in GA treated cell (Fig 3F) show that effective GA doses were used. These results indicate that the effect in NS1 secretion observed after FKBP52 silencing is not related with HSP90 activity and was rather associated with the function of the CCC. Finally, the expression of Cy40 was significantly knock down at 48h post siRNA transfection (Fig 4A). A reduction in about 60% of lipid droplets counting per cell was observed in transfected cells (Fig 4B). Yet, neither NS1 secretion nor virus release was affected by Cy40 silencing (Fig 4C and 4D). This result suggests that while Cy40 participates in cholesterol traffic it may be dispensable for NS1 secretion.

**Fig 3.**
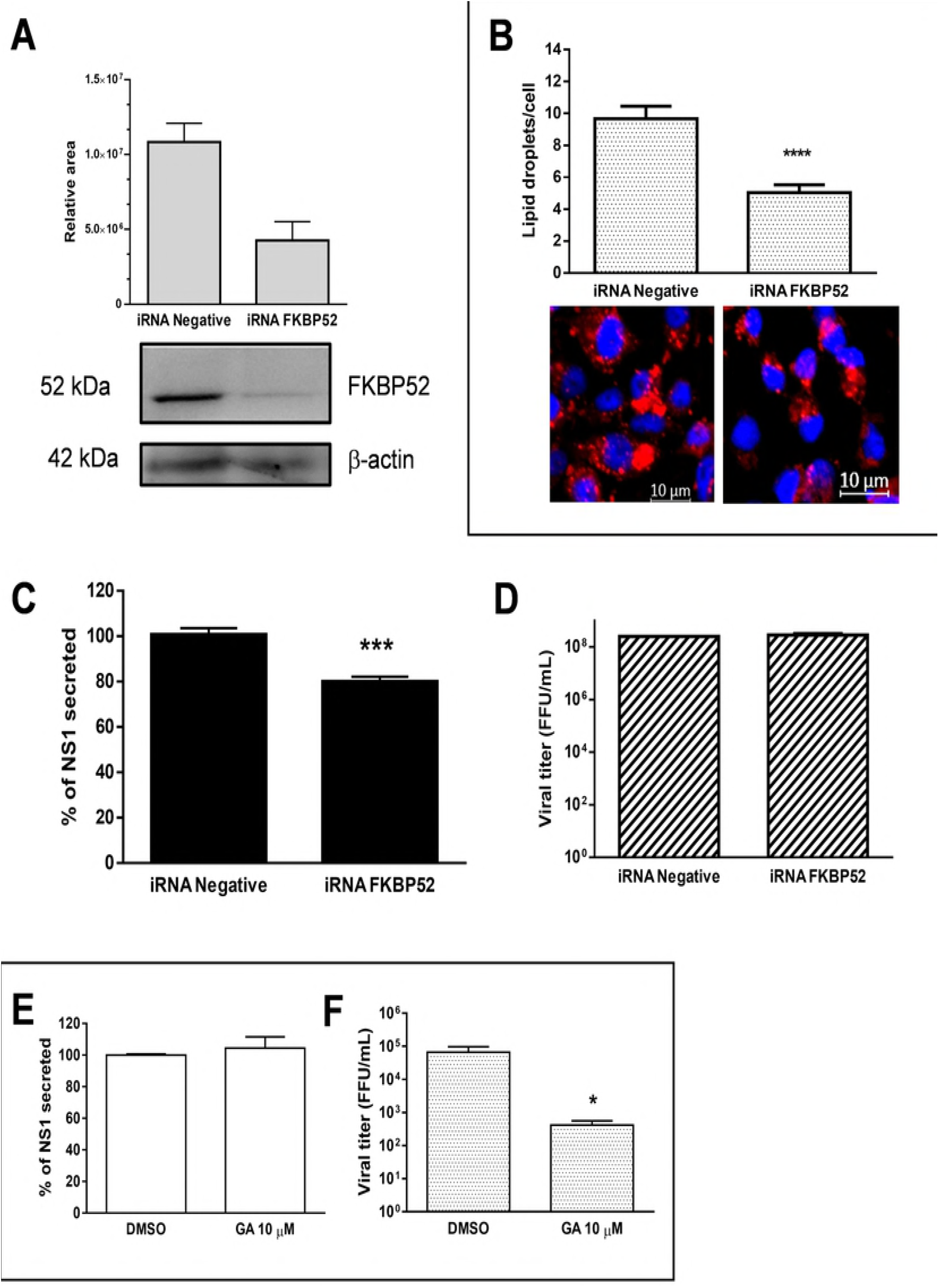
NS1 secretion is inhibited in mosquito cells transfected with FKBP52 siRNA. **A** C6/36 cells were transfected with 50nM siRNA targeting FKBP52 or with AllStars Negative Control siRNA. Protein expression was measured by western blot after 48 hours. After 24 h post transfection, cells were infected with DENV4 during 1 h, at MOI of 3. **B** LD count in C6/36 cells silenced for FKBP52 expression. LD counting in FKBP52 and negative knockdown was measured by LD staining with Nile red. LD were counted in maximum projection images in at least 20 treated cells using spot detector plugin in Icy software. LD median count differences were compared using Student’s t test denoted by * (P ≤ 0.05). **C** Secreted DENV NS1 in siRNA transfected C6/36 infected cells. NS1 was measured after 48h post transfection using Platelia NS1 Ag (Bio-Rad) and represented as % of NS1 secreted compared to negative siRNA. **D** Viral titers in supernatants were measured by focus forming assay in BHK-21 cells following standard procedures. **E** Independence of the FKBP52 co-chaperone activity on HSP90 was assayed by geldanamycin (GA) treatment. HSP90 inhibition is not responsible for reduction in NS1 secretion. C6/36 cells were treated with 10μM geldanamycin a specific inhibitor of HSP90. **F** Percentage of secreted NS1 and viral titers were measured in supernatants after 24 h of treatment. ***C***, ***D, E*** *and* ***F***; All data are mean of 3 independent experiments ± standard error (error bars), and significant differences were compared using Student’s t test denoted by * (P ≤ 0.05).

**Fig 4.**
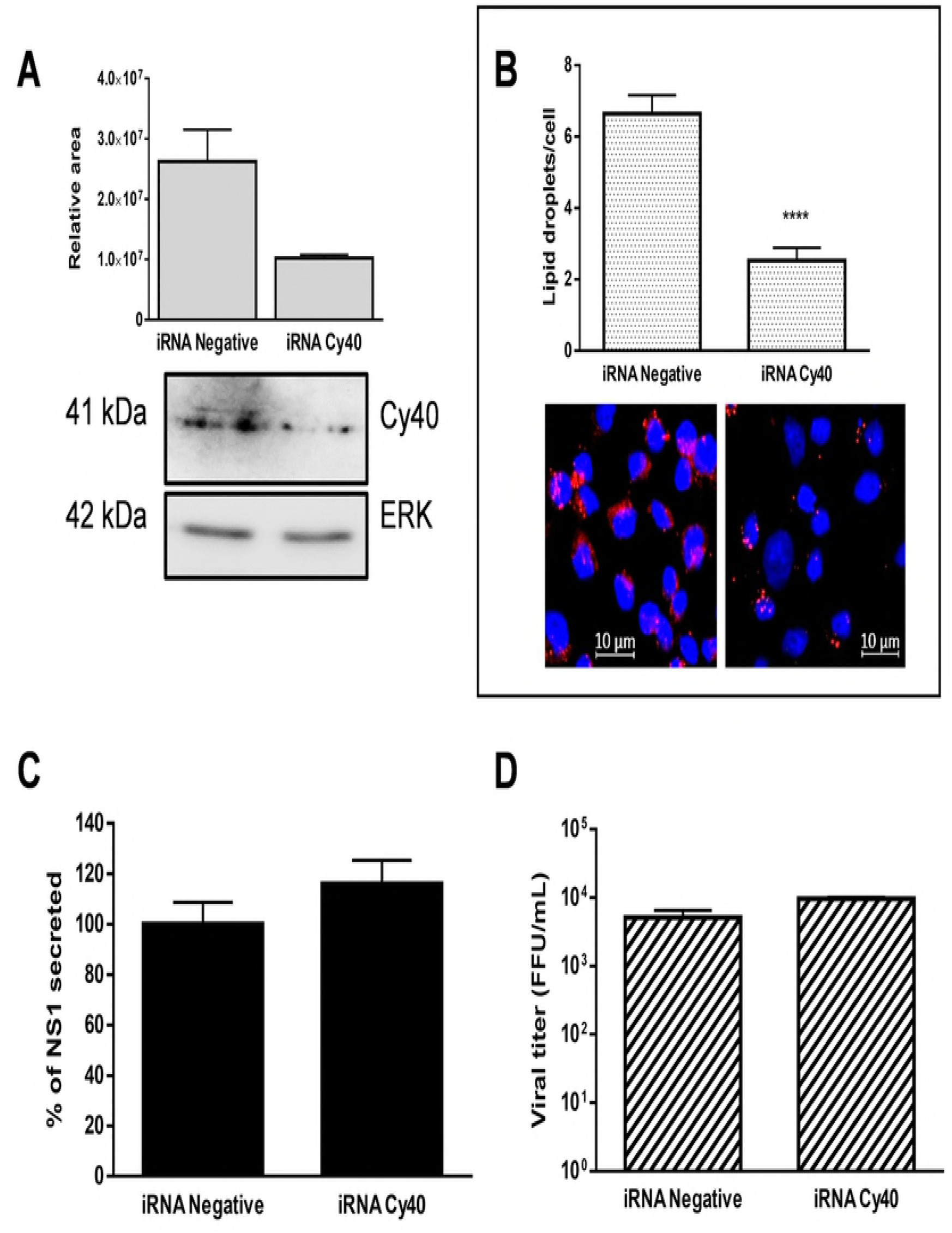
NS1 secretion is not affected in mosquito cells transfected with Cy40 siRNA. **A** C6/36 cells were transfected with 50nM siRNA targeting Cy40 or with All Stars Negative Control siRNA. Protein expression was measured by western blot after 48 hours. After 24 h post transfection, cells were infected with DENV4 during 1h at MOI of 3. **B** LD count in C6/36 cells silenced for Cy40 expression. LD counting in Cy40 and negative knockdown was measured by LD staining with Nile red. LD were counted in maximum projection images in at least 20 treated cells using spot detector plugin in Icy software. LD median count differences were compared using Student’s t test denoted by * (P ≤ 0.05). **C** Secreted DENV NS1 was measured after 48h post transfection using Platelia NS1 Ag (Bio-Rad) and represented as % of NS1 secreted compared to negative siRNA. **D** Viral titers in supernatants were measured by focus forming assay in BHK-21 cells following standard procedures. ***C****, and* ***D***; All data are mean of 3 independent experiments ± standard error, and significant differences were compared using Student’s t test denoted by * (P ≤ 0.05).

Since the hexameric NS1 is rich in cholesterol and triglycerides, a possible interaction between DENV NS1 and lipid droplets was tested through co-localization assays. DENV infection results in an increased in LD in mosquito cells (S2 Fig), yet NS1 did not show significant PCC values with LD (S4 Fig). Finally, the interference with the chaperones that conform the CCC in non-infected cell resulted in a significant reduction in the count of lipid droplets per cell in all cases. This decrease is similar to the observed in cells treated with MβCD or subjected to serum starvation (S3 Fig). These data generated in mosquito cells, endorse the role proposed for the CCC in lipid traffic and homeostasis in vertebrate cell [12,13].

### Dengue virus infection increases the expression of CCC proteins

Given the relatively high amounts of NS1 synthetized and the expected high amounts of lipids required to form the hexamer, and given the importance of CCC in NS1 traffic and lipid homeostasis, the effect of dengue virus infection in the CCC protein levels was evaluated. Western blot analysis of DENV infected C6/36 cells showed a 3.5 and 2.0-fold increase in CAV-1 and FKBP52 expression, respectively, at 24 hpi; meanwhile, no changes in the expression of Cy40 and CyA were observed (Fig 5A). The infection of BHK-21 cell run in parallel showed no changes in the expression of any of the proteins associated with the CCC (Fig. 5B). Of note, a 1.6-fold increase in CAV-1 expression levels was observed in C6/36 cell treated with MβCD for 24 h (Fig. 5C), suggesting that the increase in CAV-1 expression is a response of the mosquito cell to alterations in lipid homeostasis.

**Fig 5.**
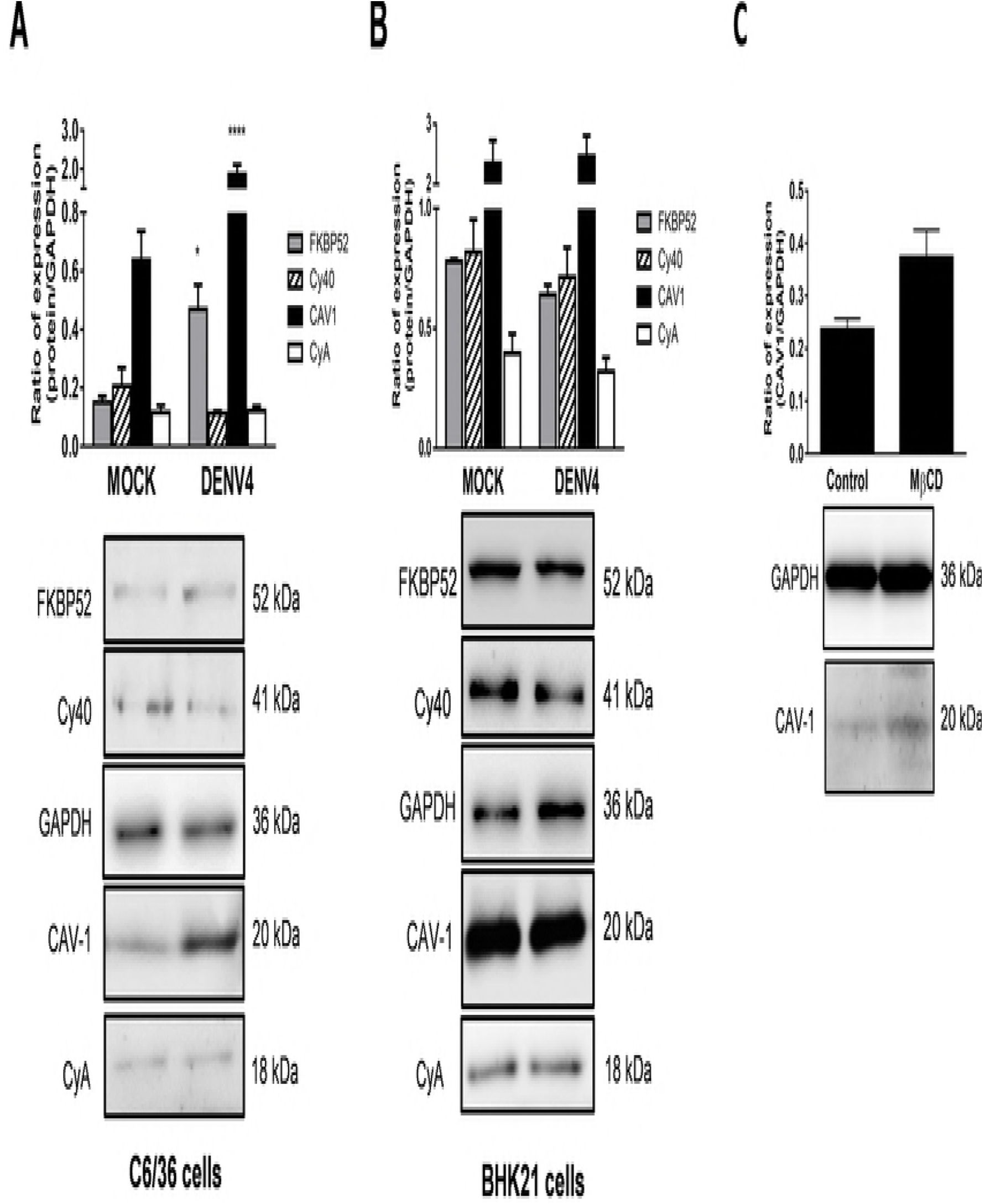
Expression of the chaperon caveolin complex is augmented in DENV infected mosquito cells. **A** Relative protein expression of CCC in DENV4-C6/36 infected cells at MOI=3. **B** Relative protein expression of CCC in DENV4-BHK-21 infected cells at MOI= 3. **C** Disruption of cholesterol flow in C6/36 cells was achieved by treatment with methyl-β-cyclodextrin (MβCD) 9 mM. CAV-1 ratio expression was determined in cells lysates by western blot and normalized with GAPDH. Values are means of 4 independent experiments ± SEM. ***A*** *and* ***B***, Cells were MOCK or DENV4 infected at MOI of 3 for 24h. The relative protein expression of caveolin chaperone complex was determined in cell lysates by the ratio of the sample value to an internal standard control (GAPDH). Ratio values are means ± SEM (n?=?4: MOCK or DENV4). Significant differences were compared using Mann–Whitney U test. Significant * (p < 0.05).

### In vertebrate cells, NS1 secretion is independent of CCC

To asses if the dependence of NS1 secretion on the CCC is exclusive of mosquito cells, we evaluated the effect of silencing of CAV-1, FKBP52 and Cy40 on NS1 secretion in infected BHK-21 cells. As shown in figure 6, the use of siRNAs significantly reduced the expression levels of the targeted proteins (Fig. 6A, 6E and 6I). Moreover, a significant reduction in lipid droplet count levels was observed after the knock down of each of the proteins of the CCC (Fig. 6D, 6H and 6L). However, no changes in NS1 secretion levels (Fig 6B, 6F and 6J) or virion release (Fig 6C, 6G, and 6K) were observed in the transfected cells. These results indicate that while the CCC plays a role in the lipid homeostasis, it plays no role in the secretion of NS1 in vertebrate cells.

**Fig 6.**
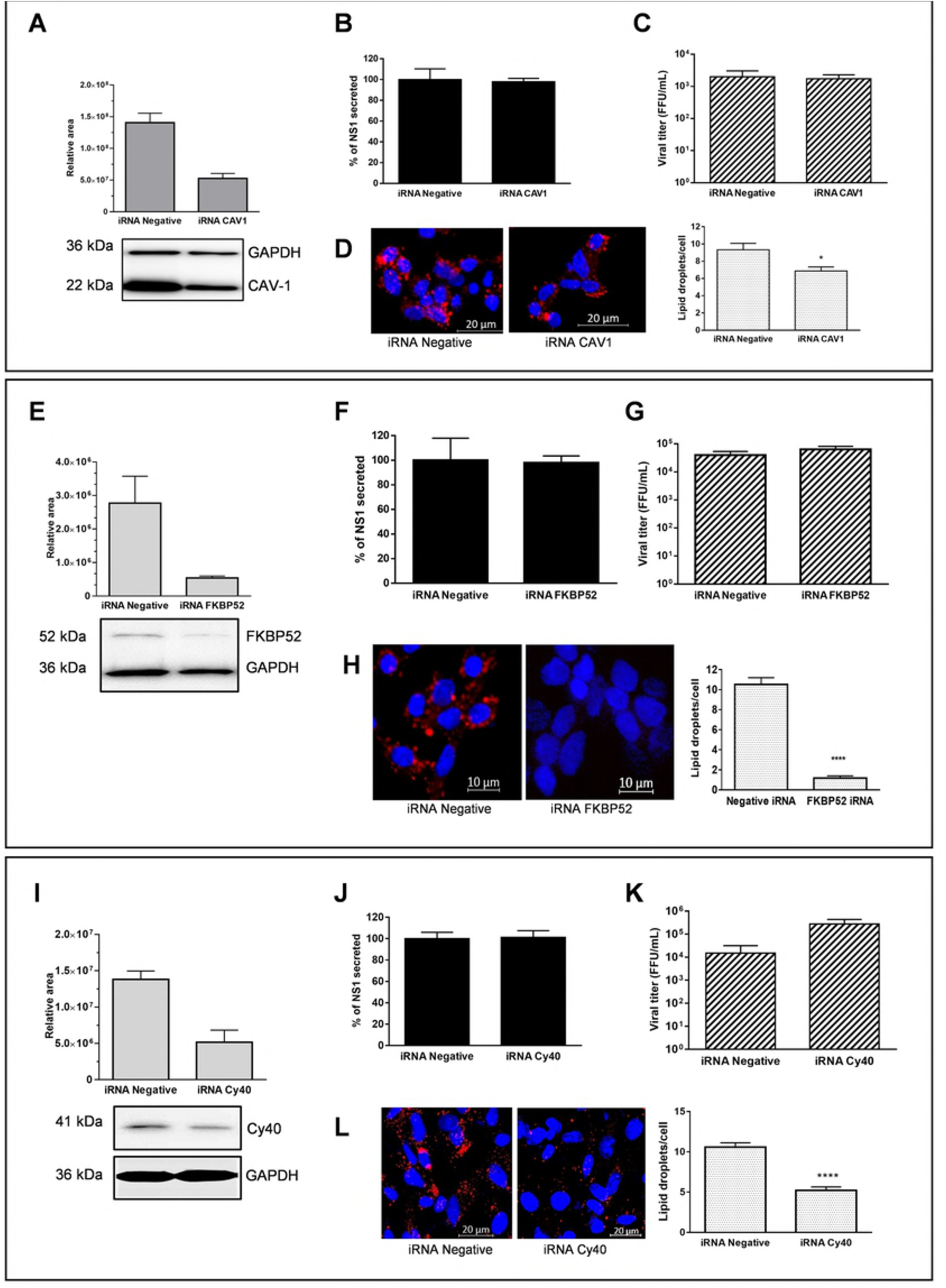
NS1 secretion is not affected in vertebrate cells transfected with CAV-1, FKBP52 or Cy40 siRNAs. **Knockdown of CAV-1. A** BHK-21 cells were transfected with 50nM siRNA targeting CAV-1 or with AllStars Negative Control siRNA. **B** Secreted DENV NS1 after CAV-1 knockdown. **C** Viral release after CAV-1 knockdown. **D** LD count in BHK-21 cells silenced for CAV-1 expression. **Knockdown of FKBP52. E** BHK-21 cells were transfected with 50nM siRNA targeting FKBP52 or with AllStars Negative Control siRNA. **F** Secreted DENV NS1 after FKBP52 knockdown. **G** Viral release after FKBP52 knockdown. **H** LD count in BHK-21 cells silenced for FKBP52 expression. **Knockdown of Cyclophilin 40. I** BHK-21 cells were transfected with 50nM siRNA targeting Cy40 or with AllStars Negative Control siRNA. **J** Secreted DENV NS1 after Cy40 knockdown. **K** Viral release after Cy40 knockdown. **L** LD count in BHK-21 cells silenced for Cy40 expression. ***A***, ***E*** *and* ***I*** Gene knockdown was assessed using Western Blotting. Protein expression was measured and normalized with GAPDH after 48 hours. After 24h post transfection, cells were infected with DENV4 at MOI of 3 for 1 h. Data are means of three experiments ± standard error of the means (SEMs). ***B***, ***F*** *and* ***J*** Secreted DENV NS1 was measured after 48h post transfection of siRNA using Platelia NS1 Ag (Bio-Rad) and represented as % of NS1 secreted compared to negative siRNA. ***C***, ***G*** *and* ***K*** Viral titers were measured by focus forming assay in BHK-21 cells. ***B***, ***C***, ***F***, ***G***, ***J*** *and* ***K***; All data are mean ± standard error of the means (SEMs) (error bars), n = 3, and significant differences were compared using Student’s t test denoted by * (P ≤ 0.05). ***D, H,*** *and* ***L***; LD was measured by LD staining with Nile red and counted in maximum projection images in at least 20 cells using spot detector plugin in Icy software. LD median count differences were compared using Student’s t test denoted by * (P ≤ 0.05).

### DENV NS1 co-localizes with the CCC in mosquito cells

Based on the dependence of NS1 secretion on the CCC observed, we evaluated the association between NS1 and CAV-1, FKBP52, Cy40 and CyA in mosquito infected C6/36 (Fig 7A, 7B, 7C and 7D) and Aag2 (Fig 7E, 7F, 7G and 7H) cells and fixed at 18 hpi. BHK-21 cells were run in parallel as controls (Fig 7I, 7J, 7K and 7L). Quantification of the co-localization levels using the Pearson Correlation Coefficient (PCC) indicated significant (PCC≥0.5) co-localization between NS1 and all the components of the CCC in both insect cell lines, especially with CAV-1 and CyA.

The co-localization between NS1 and CCC was always observed in the cytoplasm, randomly distributed and not forming any specific structure. As expected, no co-localization (PCC≤0.2) of NS1 with the proteins of the CCC was observed in BHK-21 cells. Finally, co-localization studies carried out in mock infected cells, suggest the association of CAV-1 with FKBP52, Cy40 and CyA, and thus the presence of the CCC, in both mosquito and vertebrate cell lines (Fig 7N and S4 Fig).

**Fig 7.**
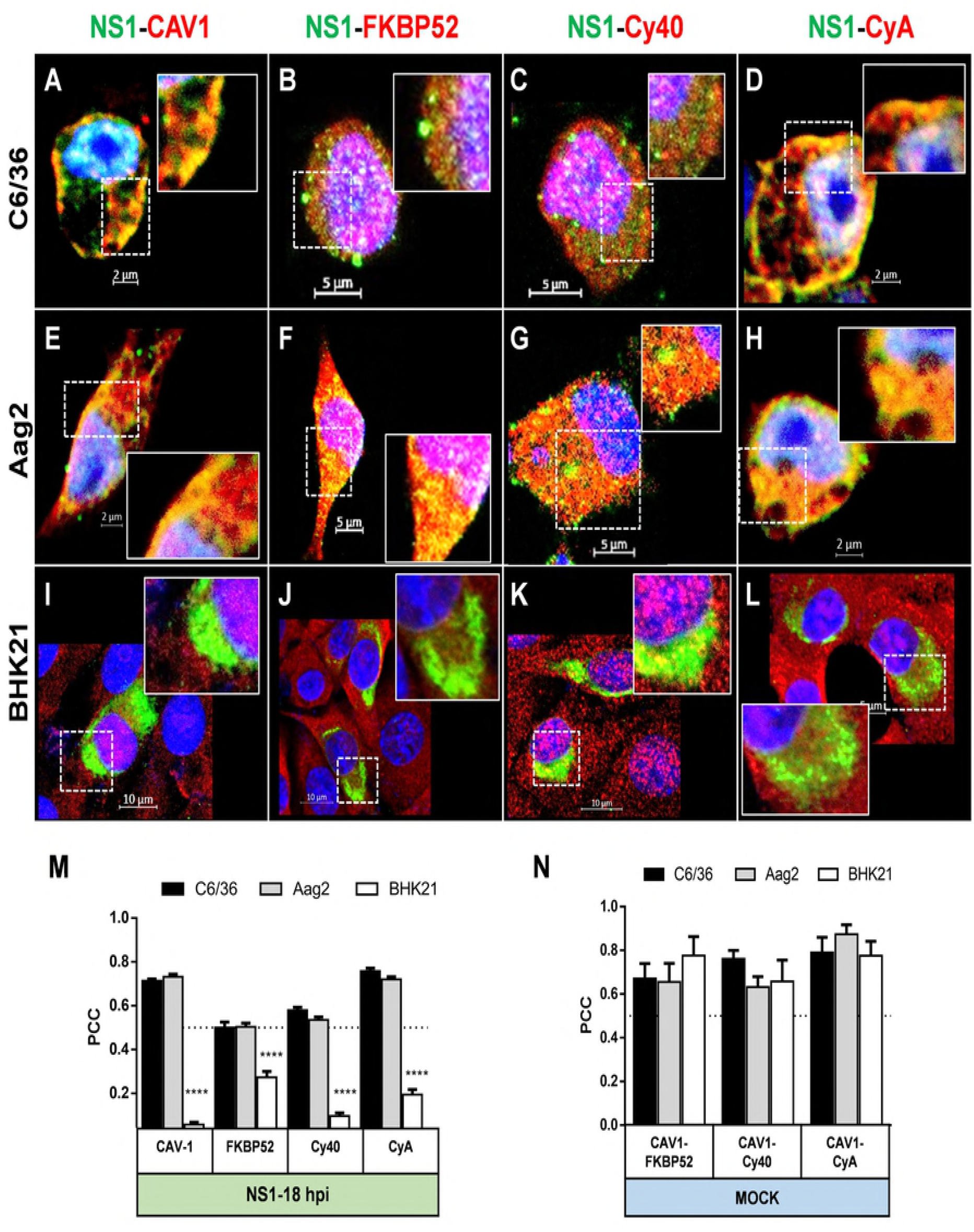
DENV NS1 co-localizes with proteins of the CCC in mosquito cells. **A**, **B, C,** and **D** Co-localization for NS1 and CCC in C6/36 cells. **E**, **F, G,** and **H** Co-localization for NS1 and CCC in Aag2 cells. **I**, **J, K,** and **L** Co-localization for NS1 and CCC in BHK-21cells. All cells were infected with DENV4 at MOI?=?3 and fixed at 18 hpi. The cells were probed against NS1 (in green), and CAV-1, FKBP52, CyA, and Cy40 (in red). Nuclei were stained with DAPI (in blue). Merged fluorescent images of DAPI, red and green channels are shown. Insert panels show magnification of the detected spots corresponding to the co-localization. The images were analyzed using a LSM 700 confocal microscope. **M** Pearson correlation coefficient (PCC) was used to evaluate degree of co-localization between NS1 and each of the CCC proteins in the 3 cell lines infected with DENV4. **N** PCC was used to evaluate degree of co-localization between CAV-1 and FKBP52, CyA and Cy40 in non-infected cells lines as control of CCC. Confocal Images of non-infected cells are shown in S4 Fig. ***M*** *and* ***N*** The images were analyzed using a LSM 700 confocal microscope with laser sections: 0.45 μm. The bars represent means ± standard error from at least 20 confocal independent cell images. Data was evaluated using the 2way ANOVA test and significant differences between groups were denoted by *(P ≤0.0001). Dotted line indicates threshold for true co-localization (PCC ≥ 0.5).

### NS1 interacts with chaperone caveolin complex in mosquito cells

The co-localization results suggest that NS1 interacts with CAV-1 and the chaperones of the CCC in DENV infected mosquito cells. To test this possibility, co-inmunoprecipitation (Co-IP) experiments were carried out using C6/36 infected cell lysates harvested at 18 hpi and mock infected cells lysates as controls (Figure 8A). Immunoprecipitation were carried using both anti-NS1 and anti-CAV-1 as primary antibodies. Then, the presence of NS1, FKBP52, Cav-1, Cy40 and CyA in the precipitated immunocomplexes was revealed by western blot. The presence of NS1 was clearly observed in the Co-IP with anti-CAV-1, indicating the interaction between NS1 and CAV-1 in infected mosquito cells. This interaction was confirmed with the detection of CAV-1 in the Co-IP with anti-NS1. In addition, the presence of FKBP52, Cy40 and CyA was also detected in the Co-IP with anti-NS1. No proteins were detected in the Co-IP with anti-NS1 when in mock infected cell lysates were used. These results taken together indicate that NS1 interacts with the CCC in DENV infected C6/36 cells. In addition, the detection of FKBP52, CyA and Cy40 in the Co-IP with anti-CAV-1 in both mock and infected cell lysates, corroborates the co-localization data indicating the existence of the CCC in mosquito cells.

**Fig 8.**
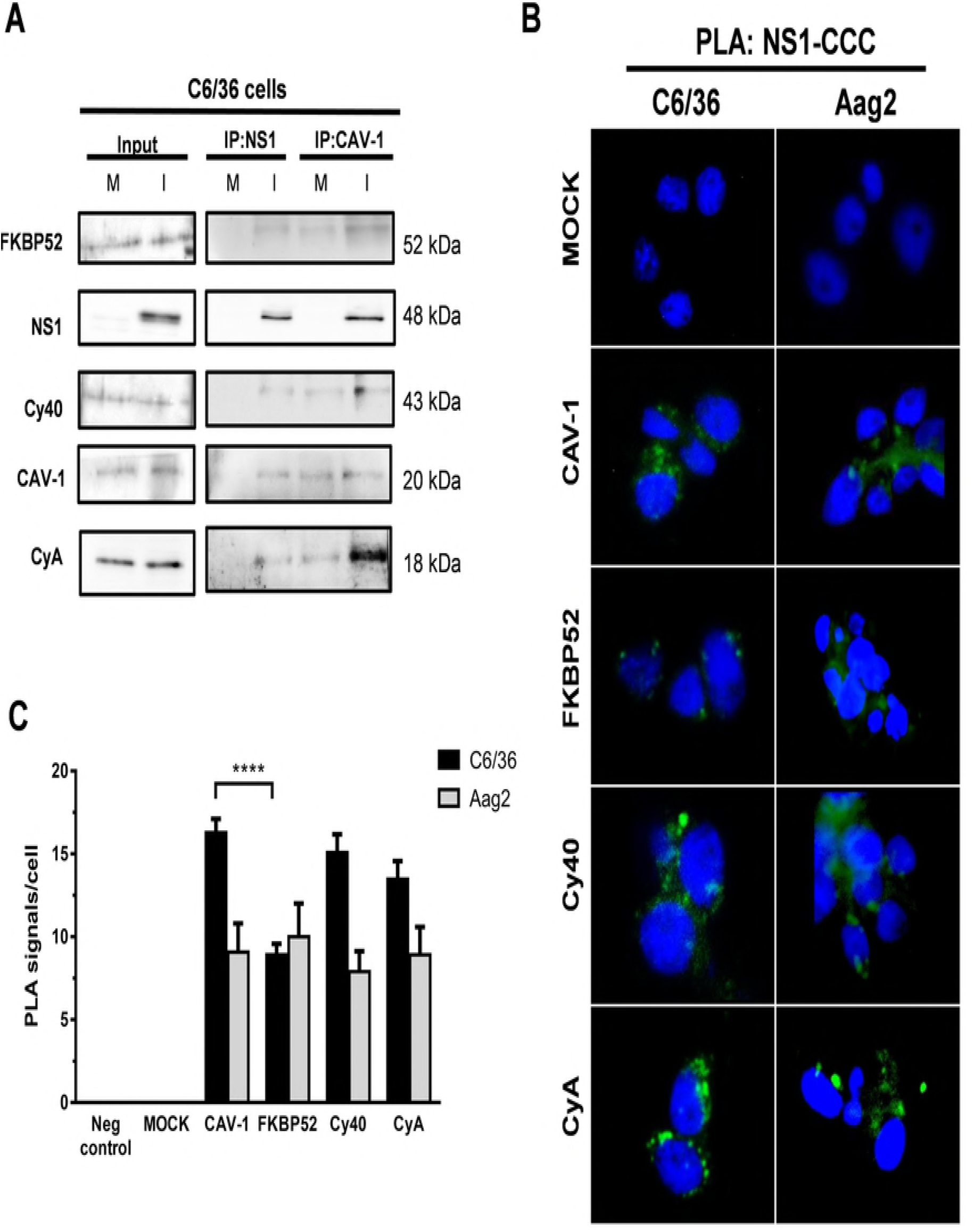
DENV NS1 interacts with proteins of the CCC in mosquito cells. **A** Immunoprecipitations assays with anti-NS1 and anti-CAV-1 antibodies in C6/36 cells. Mock (M) and DENV4-infected (I) cell lysates (18hpi) were processed for immunoprecipitation (IP) using anti-CAV-1 polyclonal antibody (n-20, Santa Cruz) and anti-NS1 polyclonal antibody (GTX124280-Genetex). After extensive washing, eluted protein complexes were analyzed for the presence of CAV-1, NS1, FKBP52, Cy40 and CyA by western-blot. Input: Mock and DENV4-infected cell lysates prior to the immunoprecipitation served as controls for protein detection. **B** Proximity ligation assay (PLA-Duolink^®^) between NS1 and the CCC in infected mosquito cells. PLA signals (green) represent dual recognition PLA against NS1 and CAV-1, FKBP52, Cy40 or CyA. Nuclei were stained with DAPI (blue). **C** Quantification of PLA signals per cell on DENV4 infected C6/36 and Aag2 cells (PLA signals greater than 3 pixels). C6/36 and Aag2 mock infected cells were incubated with both primary antibodies, and infected cells incubated without CAV-1 antibody, were included as negative controls. The bars represent means ± standard error from at least 20 cells. Data was evaluated using the Student t test and significant differences of CCC against CAV-1 were denoted by *(P ≤0.05).

To confirm direct interaction between NS1 and proteins of CCC, we performed a proximity ligation assay (PLA) analysis in DENV infected C6/36 and Aag2 infected cells. The PLA allows the detection of protein-protein interactions, while preserving the cell architecture and in addition the generated signal can be quantified. In agreement with previous results [10], a positive signal (green dots) was detected in both DENV-infected mosquito cells stained for NS1 and CAV-1, confirming that NS1 and CAV-1 directly interact in infected mosquito cells (Fig 8B). In addition, positive signals were detected between NS1 and each of the chaperone proteins of CCC; Cy40, FKBP52, and CyA (Fig 8B), indicating that NS1 interacts with the whole CCC in both cell lines. In all cases, the PLA signals were located in the cytoplasm, in agreement with the co-localization results shown in Figure 7. Quantification of the PLA signal per cell suggests that NS1 interacts equivalently with all the components of the CCC, although significantly less signal was observed for FKBP52 and NS1 in C6/36 cells. No fluorescent spots were detected in mock-infected cells incubated with both primary antibodies or in infected cells incubated with anti-NS1 in the absence of anti-CAV-1, signifying the specificity of the signal (Figure 8B and 8C).

### ZIKV NS1 follows an CAV-1-dependent unconventional secretion in mosquito cells

NS1 is the only viral non-structural protein secreted from DENV-infected cells. Zika virus (ZIKV) secretes NS1 efficiently from infected cells. Other *Flaviviruses* like Yellow Fever virus (YFV) also secrete NS1 albeit in very low amounts compared with DENV or ZIKV [24]. In order to compare with DENV, we tested the secretion properties of ZIKV and YFV NS1 in infected C6/36 and Vero-E6 cells. The presence of ZIKV NS1 was detected by ELISA in the cell supernatants of both cell lines at 48 hpi, whereas YFV NS1 was detected at 72 hpi, by western blot assay (Fig 9A). Vero-E6 cells were used as vertebrate cells in these experiments since very low levels of infection were detected in BHK-21 cells with ZIKV and YFV (data not shown). The secretion of YFV NS1 was found to be sensitive to BFA treatment in both mosquito and vertebrate cells (Fig. 9A). In contrast, the secretion of ZIKV NS1 was found to be insensitive to BFA and GCA treatment in infected mosquito, but not in infected vertebrate cells (Fig 9B). In addition, the secretion of ZIKV NS1 was reduced in mosquito, but not in vertebrate cells, treated with the CyA inhibitor CsA (Fig 9C). Co-localizations experiments carried out in C6/36, Aag2 and Vero-E6 cells infected with ZIKV and YFV, showed a stronger, and a significantly different, co-localization between ZIKV NS1 and CAV-1 than between YFV NS1 and CAV-1, in both mosquito cell lines. No co-localization of ZIKV or YFV NS1 with CAV-1 was observed in vertebrate cell line (Fig 9D and 9E). Of note, the co-localization data support the predictions derived from the presence of a conserved caveolin binding (CBD) domain in ZIKV (and DENV) but not in YFV NS1 (Fig 9F). Finally, the secretion of ZIKV and YFV virions was affected by the BFA and GCA treatment in both mosquito and vertebrate cells (S5 Fig). All the data taken together suggest that, just like DENV NS1, ZIKV NS1 is secreted from mosquito cells, following an unconventional secretory route in association with CAV-1 and the CCC, while a classical secretory route is followed for ZIKV NS1 secretion in vertebrate cells. On the other hand, YFV NS1 seems to be secreted by a classical secretory pathway in both cells types.

**Fig 9.**
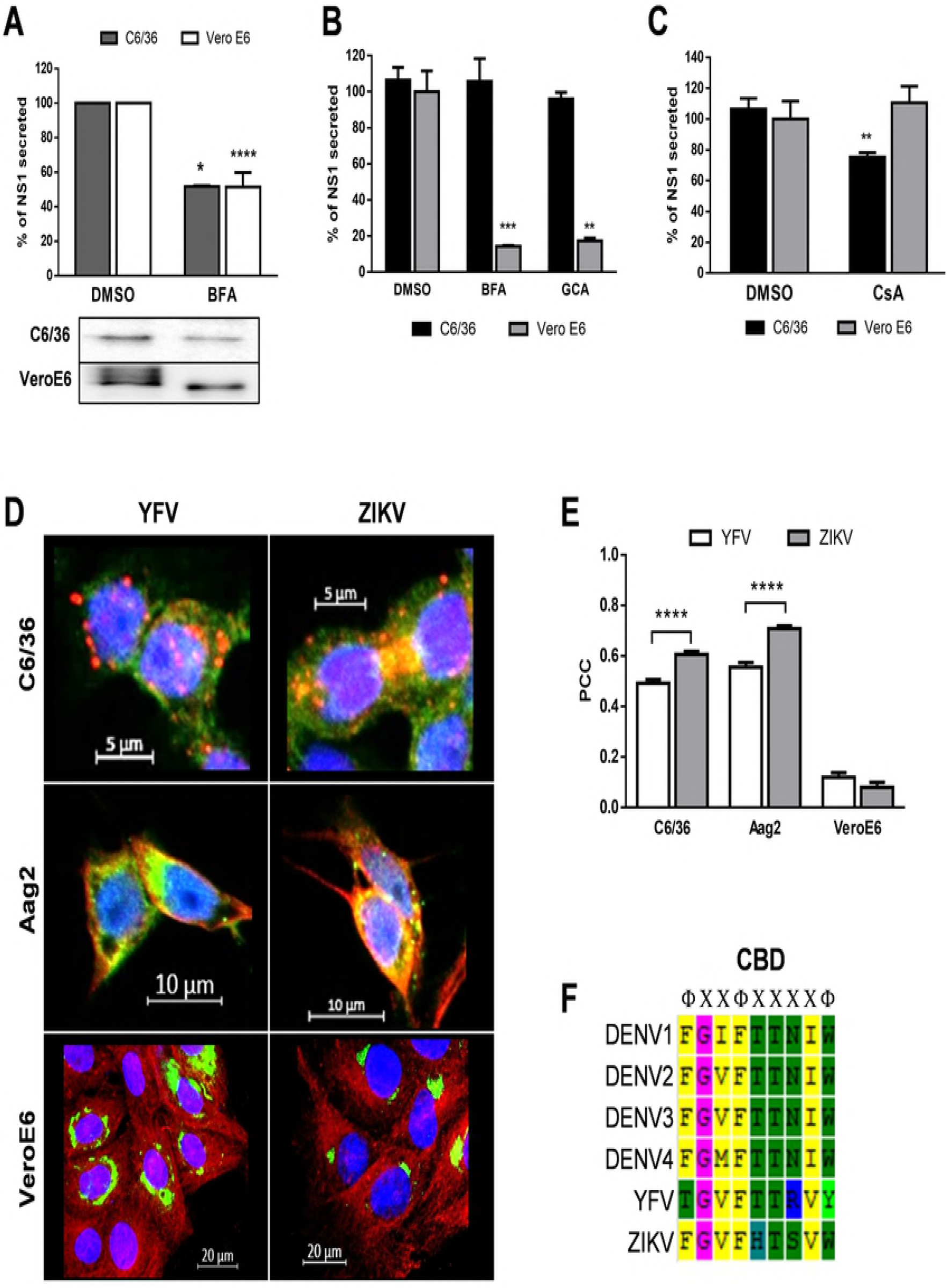
ZIKV and YFV NS1 secretion in infected mosquito cells. **A** Inhibition of classical secretory pathway in YFV infected C6/36 and Vero E6 cells at 72hpi. **B** Inhibition of classical secretory pathway in ZIKV infected C6/36 and Vero E6 cells at 48hpi. **C** ZIKV NS1 secretion is reduced in C6/36 cells treated with CsA 9 μM. **D** Co-localization between YFV NS1 or ZIKV NS1 and CAV-1 in mosquito and VeroE6 cells. C6/36, Aag2 and VeroE6 cells were infected at MOI of 1 for 48 h. Cells were probed for CAV-1 (shown in red), viral NS1 (shown in green) and nuclei stained with DAPI (blue). **E** Pearson correlation coefficients (PCC) for CAV1-NS1 were measured in at least 20 confocal independent images with 0.47 μM laser sections. The bars represent means ± standard error. Data was evaluated using the 2way ANOVA test and significant differences are denoted by *(P ≤0.05). **F** Protein sequence alignment of caveolin binding domain (CBD) in *Flavivirus* NS1. Symbols: Φ, aromatic amino acid (F/Y/W) and X, any amino acid. ***A***, ***B*** *and* ***C***; Cells were infected with YFV or ZIKV at MOI of 1 for 1 h. The cells were washed 3 times and after treated with DMSO (control), 7μM BFA, 27μM GCA or 9μM CsA. For ZIKV infection supernatants were harvested at 48 hpi and for YFV infection supernatants were collected 72 hpi. Secreted ZIKV NS1 was measured using Platelia NS1 Ag (Bio-Rad) and represented as percentage of secreted NS1 compared to DMSO. Secreted YFV NS1 was evaluated by western blot. The experiments were performed in triplicate, and the bars represent means ± standard error. Data was evaluated using the 2way ANOVA test and significant differences compared with DMSO treatment group, respectively, are denoted by *(P ≤0.05).

### Modeling of molecular interactions between DENV NS1 and the chaperone caveolin complex

To better understand the molecular interactions between the dimeric DENV NS1 protein and the CCC, we constructed a human CAV-1 3D model using RaptorX server (http://raptorx.uchicago.edu) [25]. The best template for the 3D CAV-1 model was 2A65 (DOI: 10.2210/pdb2a65/pdb) with a p-value 2.69e-03. Molecular docking predicts a favorable interaction between the NS1 hydrophobic domain including the grease fingers, where the CBD is located, with a pocket in the CAV-1 scaffolding domain (CSD) with distances less 8Ǻ (Fig 9A and 9B). The molecular docking results obtained with CAV-1 and NS1 were used as a control to carry on additional docking experiments between the DENV NS1 or CAV-1 and the CCC chaperones FKBP52, Cy40 and CyA. Calculations of the energy binding of several docking clusters showed that the DENV NS1 has the lowest and more favorable energy for CAV-1, compared to the energy binding obtained for FKBP52, Cy40 or CyA. In addition, lower energies were obtained for the binding of CAV-1 with the other chaperones of the CCC, than for the binding of NS1 with those same chaperones (Fig 9C). Based in the free energies data, we predict a model of interaction of NS1 with the CCC proteins in which FKBP52, Cy40 and CyA are directly bound to CAV-1, and NS1 is bound to the CCC through the CAV-1 scaffolding domain (Fig 9D).

## DISCUSSION

Our previous work demonstrated that DENV NS1 secretion in mosquito cell lines follows a BFA-insensitivity secretion pathway that depends on CAV-1; moreover, NS1 was found to interact directly with CAV-1 [10]. Therefore, in the present work, we explored the association of NS1 with proteins involved, together with CAV-1, in the intracellular transport of cholesterol and the dependence of NS1 secretion on such association. The results revealed that NS1 is associated to the chaperone caveolin complex (CAV-1, Cy40, FKBP52 and CyA) and that the integrity of this complex is necessary for the secretion of NS1 in mosquito infected cells. Additional data suggest that in virus infected mosquito cells, ZIKV NS1, but not yellow fever virus NS1, also associates with CAV-1 and usurps the cholesterol traffic pathways for secretion. Thus, the lipid content or molecular structure of the DENV NS1 seems to have taken advantage of mosquito cell cholesterol transport to guarantee the efficient NS1 secretion in the mosquito vector. Interestingly, even though the CCC is present in vertebrate cells, NS1 is not associated to it and follows a strict classical secretion pathway.

The data obtained with the earlier inhibitors of the classical secretory pathway, GCA and Fli-06, corroborate and expand the previous data obtained with BFA, regarding the use of NS1 of non-classical secretory pathways in mosquito cells. GCA inhibits the COPI-II recruitment and ER exit site (ERES) transport to Golgi [20]. GCA-insensitivity indicates that the NS1 and CAV-1 interaction initiates in the ER compartment. In agreement, treatment of mosquito cells with Fli-06 showed no effect on NS1 secretion, indicating that the interaction between NS1 and CCC is taken place very early after NS1 synthesis, before the viral protein is transported to the ERES [21,22].

Few works have reported a luciferase secretion system in insect cell lines [26]. Yet, as an additional tool to study classical secretion in mosquito cell lines, an efficient *Gaussia* luciferase reporter system that enables the detection of Luciferase in cell supernatants 6 hours after transfection was developed. The secretion of luciferase was significantly affected after treatment with GCA and BFA (data not shown); these data, together with the observed inhibition of viral particle release, indicates the existence of a robust classical secretion pathway in mosquito cells. Although, mosquito cells present both classical and unconventional secretory pathways, DENV NS1 is secreted bypassing the Golgi apparatus. A study in mosquito cells demonstrated that insect cells have an active ER-retention protein system; the KDEL c-terminal peptide functions as a ER-retention signal for some proteins [27]. NS1 lacks a KDEL signal, yet it presents a conserved and well exposed caveolin biding domain (CBD) [10]. Thus, NS1 unconventional secretion in mosquito cells is likely associated to this CAV-1 motif. However, the CBD may not be the unique factor to determine unconventional secretion. A role for amino acids at position 10 and 11 in flavivirus NS1 secretion in mosquito cells have been proposed [28].

The CCC was first described in vertebrate cells as a non-vesicular cholesterol transport within the cell [14]. Yet, to our knowledge there are no previous reports of the presence of the CCC in any insect or mosquito cell line. Thus, given that there are differences in the lipid homeostasis in mosquito cells, including the absence de novo cholesterol synthesis pathways, it became necessary to demonstrate the existence of the CCC in mosquito cells and to show a role for the complex in lipid homeostasis, as have been shown for vertebrate cells [12,13]. Co-localization and immunoprecipitation experiments indicated that in uninfected mosquito cells, CAV-1 is indeed associated with all 3 chaperones proposed to conform the complex; FKPB52, Cy40 and CyA; moreover, knocking down the expression of CAV-1 (data not shown) or any of the associated chaperones resulted in a significant decrease in the lipid droplet count per cell, indicating also a role for the CCC in lipid homeostasis in mosquito cells. In agreement with our results, ER chaperones have been reported necessary for the secretion of recombinant proteins expressed in insect cells using a baculovirus system [29,30].

The NS1 and CCC interactions in infected mosquito cells was validated by colocalization, co-inmunoprecipitation and proximity ligation assays. Firstly, the co-localization analysis and co-inmunoprecipitation assays carried out in mock infected cells assays demonstrated direct interaction of FKBP52, Cy40 and CyA with CAV-1, again supporting the existence of the CCC in mosquito cells. In agreement with our previous report, strong CAV-1 and NS1 interaction were detected by both techniques and used as reference for the new analysis with the CCC. The capture with anti-NS1 antibodies indicated that FKBP52, Cy40 and CyA interact with NS1 in infected cells. These results were confirmed by proximity ligation assays, a powerful approach which allows to determine and quantify protein-protein interactions in situ [31]. PLA signals between NS1 and each of the CCC proteins in mosquito infected cells demonstrated be similar to the observed between CAV-1 and FKBP52, Cy40 and CyA [10]. CAV-1 dependent secretion have been reported for non-viral proteins such as Kallikrein 6 in Colon Cancer Cells and Cyr61 in lung cells [32,33] as well as non-structural viral proteins such as rotavirus NSP4 [34,35].

The structure of the CCC is unknown; and crystal models exist all 3 chaperones but not for caveolin-1. We performed docking molecular analysis with the dimeric NS1 and complete CAV-1 (3D modeled in this work), in addition with the CCC proteins. An approximated *in silico* model for NS1 and CAV-1 interaction showed very high favorable energy biding and suggest interaction through the plasma membrane [10]. The molecular docking also allows measuring the energy binding, and distances between both crystals. The CBD in NS1 interacts with scaffolding domain (CSD) with a distance of 5Ǻ approximately. Further analysis also showed, high energy binding through hydrophobic dynamics, between the NS1 β-roll domain and FKBP52, CyA and Cy40 with distances under 6Ǻ. However, the model predicts all these proteins interacting at the same domain in NS1 (data not shown), which seems unlikely, due to steric effects. Thus, we favor an alternative molecular docking model, based on interaction energies, in which NS1 is bound directly to CAV-1, and in turn FKBP52, CyA and Cy40 are bound to CAV-1 (Fig. 10D). This model still allowed interaction of NS1 with the CCC through membranes. However, if the NS1-CCC complex will traffic to the plasma membrane as a free-cytosolic complex or membrane associated is totally unknown.

**Fig 10.**
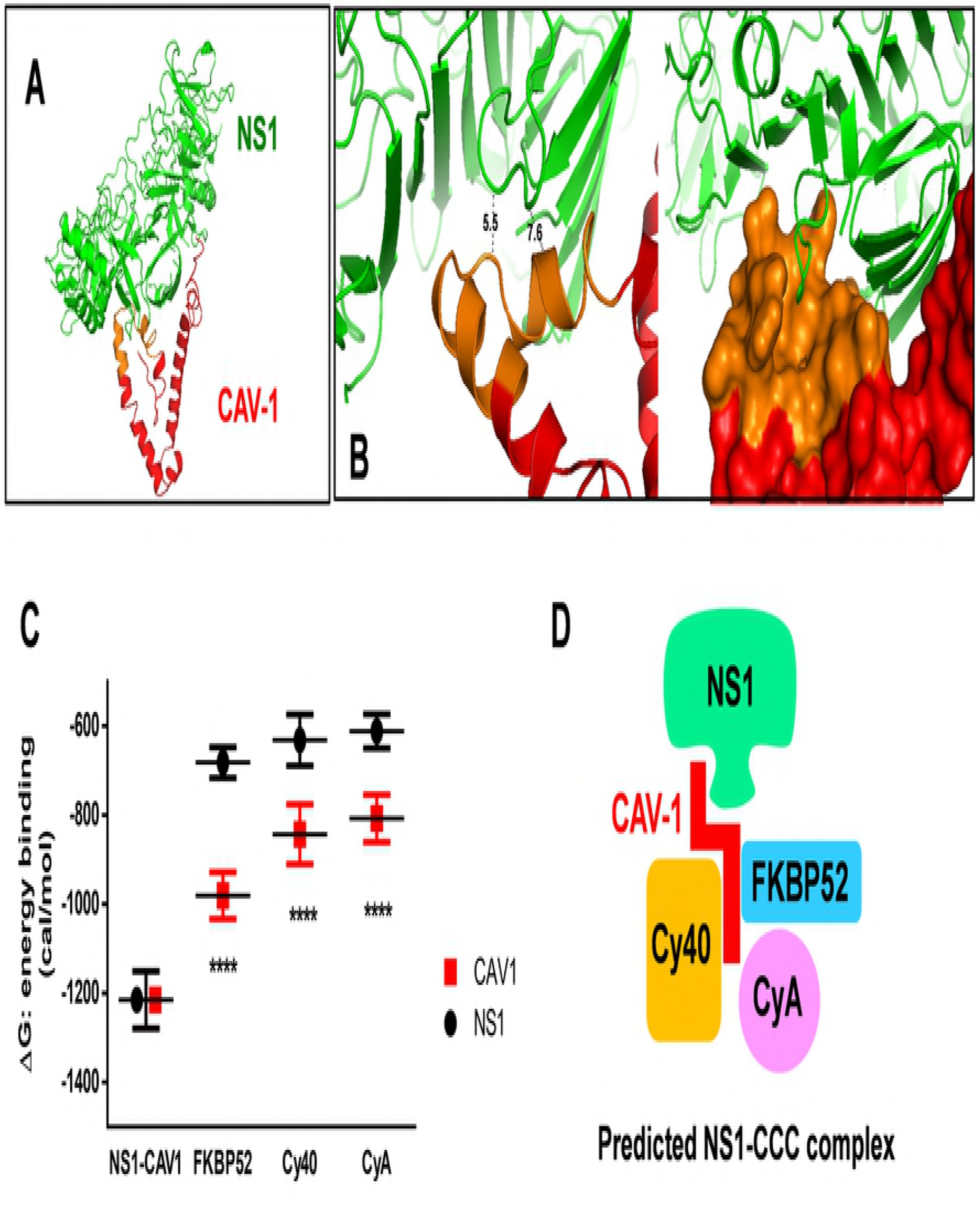
*In silico* prediction of the interaction of DENV NS1 with the CCC. **A** Interaction of NS1 and 3D model of human CAV-1 (isoform alpha, NP_001744.2). CAV-1 model (red) was retrieved from RaptorX server [25]. CAV-1 scaffolding domain (CSD), indicated in orange, dimeric DENV2 NS1 (PDB:4O6B) in green. **B** Interface distances in docking simulation of DENV NS1 and CAV-1. Dotted lines show measurement distances between aminoacids in Å as estimated by The PyMOL Molecular Graphics System (Version 2.0 Schrödinger, LLC). **C** Intermolecular binding energy for DENV NS1 and the CCC proteins. Energies of twenty clusters of protein dockings were plotted. ΔG (cal/mol) were estimated by ClusPro server (https://cluspro.bu.edu/home.php). **D** Modeled prediction of DENV NS1 and CCC based on the intermolecular binding energies.

Dependence of NS1 secretion on the integrity of the CCC was evaluated using gene knockdown for FKBP52 and Cy40 or drug inhibition for CyA. Again, the observed reduction in the number of lipid droplets per cell after each treatment, similar to the decrease observed after serum starvation and Methyl-β-Cyclodextrin (MβCD) treatment (S3 Fig), support a role for the CCC in the lipid homeostasis in mosquito cells. FKBP52 knockdown decreases NS1 secretion while viral release was not affected. HSP90 has been associated to viral replication of chikungunya virus, *Arteriviridae* viruses, Hepatitis C virus and minor effect on DENV replication [36–39], but we could demonstrate the association of this co-chaperone of HSP90 to viral protein trafficking. CyA inhibition by cell treatment with CsA also resulted in diminished NS1 secretion in mosquito cells, without affecting viral release. However, CyA has been associated to viral packing and viral release in other viruses as coronaviruses, Hepatitis C virus, HIV, and others [40–42]. Finally, Cy40 knockdown had not effect in either NS1 secretion or viral release. This result suggests a dispensable function of Cy40 in the secretion of NS1 in mosquito cell; yet, this is an unexpected result for which we have no obvious explanation, given the observed association between NS1 and Cy40, which appears to be indeed part of the CCC. Cy40 may also have an unknown regulatory activity in the CCC complex, becoming a dispensable component for the NS1 traffic in the mosquito infected cells. However, Cy40 mitochondrial function is essential for viral fitness in human coronaviruses and HCV [43,44]. Finally and regardless, the previous data indicating no interaction of the CCC proteins with NS1 in infected vertebrate cells, we wanted to explore the effect of FKBP52 and Cy40 knockdown and CyA inhibition in BHK-21 DENV infected cells. As expected, alteration in the levels or function of these proteins resulted in a significant decrease in the number of lipid droplets but had no effect on NS1 secretion, indicating that the CCC plays a role in lipid homeostasis in vertebrate cells but have no role in the secretion of NS1 and supporting a classical secretion pathway for NS1 in vertebrate cells.

Profound changes in lipid homeostasis have been observed in DENV infected mosquito cells [45] which may in part respond, to the demand of high amounts of cholesterol required for the hexameric NS1 secretion [6]. Interestingly, the levels of CAV-1 and FKBP52 were found to increase during DENV infection in mosquito, but not in vertebrate, cells. Mosquitos are cholesterol; thus, CAV-1 and FKBP52 increased expression may result as a response to increments in cholesterol demand upon infection [46,47]. As pointed out by the increased in CAV-1 expression observed after MBCD treatment of mock infected cells.

NS1 lipid cargo indicates high requirements of cholesterol and triglycerides in infected cells [6]. In hepatic cells, DENV infection induces an increase in the activity of cholesterol biosynthesis enzymes [48]. The results showing lipid droplet decay during CCC inhibition in mosquito cells, suggest a possible association between NS1 and LD during its traffic. To explore this possibility, co-localization parameters between LD and NS1 in mosquito infected cells were measured (S2 Fig). However, very low value for true co-localization between NS1 and LD particle were observed in both infected mosquito cell lines. Lipids droplets are reservoir of triglycerides and cholesterol for viral production. In addition, the C capsid protein has been shown to colocalize with LD in infected cells and this interaction is necessary for genome encapsidation [49]. In general flaviviruses increase production of free fatty acids, and cholesterol to favor assembly of viral particles [50] and NS1 secretion is affected in cells where lipid metabolism is altered, yet LD do not appear to participate in the traffic of this viral protein.

Previous results indicated that the interaction CAV-1 and NS1 is important for the secretion of NS1 in infected mosquito cells [10]. NS1 presents a CBD (ΦXXΦXxXXΦ, where Φ is an aromatic residue and X any amino acid) that is well conserved among the 4 DENV serotypes and is also conserved in the ZIKV, but not in the YFV NS1, where a T to F substitution is found in the first aromatic residue. Since changes in the aromatic residues of the CBD are important in the interaction with CAV-1 [51], the possible interactions of the NS1 of ZIKV and YFV with CAV-1 and the role of such interactions in influencing the secretory routes of NS1 from infected mosquito cells, were evaluated. The colocalization results obtained in ZIKV and YFV infected cells and well as the differential effect of BFA and GCA treatment in the secretion of NS1 of both viruses, taken together support the notion that indeed, in mosquito cells, but not in vertebrate cells, the presence of an intact CBD is necessary for the interaction between NS1 and CAV-1, and that such interaction is necessary to drive to NS1 to be secreted by a non-classical secretory route, in association with the CCC. YFV NS1 showed a favored classical Golgi secretion pathway, while ZIKV NS1 secretion is following an unconventional secretion as we have been described for DENV. Thus, the CBD appears to be a molecular determinant necessary to direct the interaction of NS1 to CAV-1 and facilitate the unconventional secretion using the cholesterol pathway and CCC [51]. Remarkably, the CBD in NS1 sequence by itself is not the only determinant to direct what type of secretion is going to take the NS1 in the cells since in vertebrate cells, NS1 from DENV and ZIKV shows no association with the CCC, and secretion is Golgi dependent. Several factors acting in concert may be necessary to keep the NS1 from reaching the ER-exit sites and entering the classical secretory pathway in mosquito cells; i) mosquito cells could be defective in a protein necessary to direct NS1 to ER-Golgi intermediate and then follow classical secretion [27]; as is suggested by the insensitivity of NS1 secretion to FLI-06 and GCA treatment, and ii) the affinity of DENV NS1 for mosquito CAV-1 may be higher for the CAV-1 of vertebrate cells; as is indeed suggested by the predicted interaction energies for both proteins where DENV NS1 presents stronger interaction for invertebrate CAV-1 than for human CAV-1 (data not shown). Figure 11 presents a schematic model for DENV NS1 secretion in infected mosquito cells.

**Fig 11.**
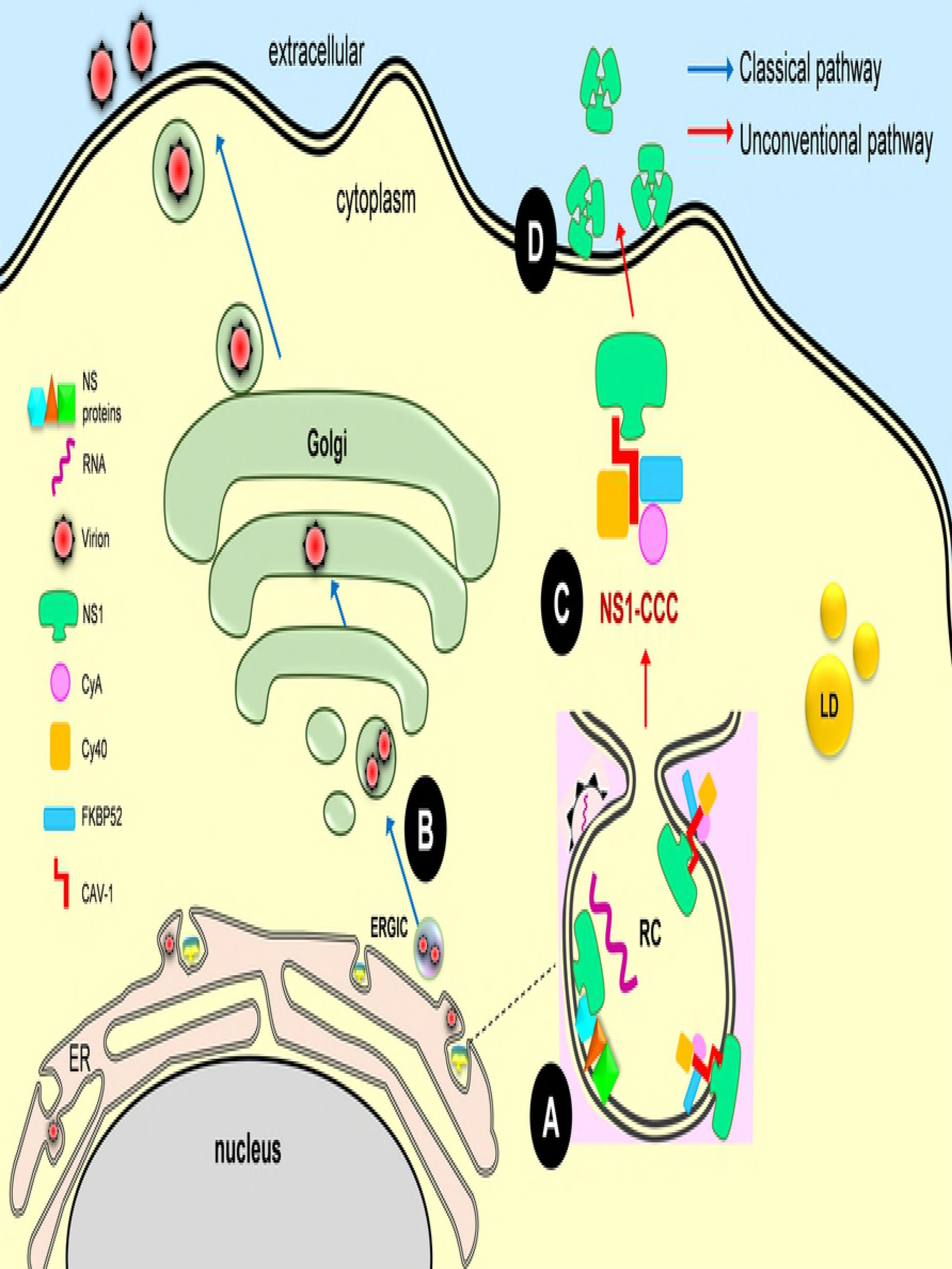
Proposed model for unconventional secretion of DENV NS1 through CCC in mosquito cells. **A** DENV replication complexes. Viral RNA is synthetized within RC with the assistance of viral non-structural proteins. NS1 interacts with the non-structural proteins to assist membrane bending and envelopment of nucleocapsids. Virions are packed and transported to ERGIC vesicles. NS1 dimer interacts with ER inner membrane in RC. **B** Virions are incorporated into vesicles within Golgi and transported through trans-Golgi network. Finally, mature virions are released to extracellular space following a Golgi-classical pathway (BFA or GCA-sensitive). **C** NS1 interacts with CAV-1 scaffolding domain through its β-roll domain in the ER-lumen. Proteins FKBP52, Cy40 and CyA interact with CAV-1 and assemble the caveolin chaperone complex (CCC). The NS1-CCC complex is detached from ER-exterior membrane by unknown mechanisms. **D** Cytoplasmic NS1-CCC complex follows the intracellular cholesterol pathway until grasp inner plasmatic cell membrane and then, hexameric NS1 is released to extracellular space. NS1 does not interact with LD during traffic. RC: replication complex; NS proteins: viral non-structural proteins; CCC: chaperone caveolin complex; ERGIC: ER-Golgi intermediate compartment; LD: lipid droplet and ER: endoplasmic reticulum.

In summary, these results indicate that in mosquito cells, DENV NS1 utilizes the CCC as part of the cholesterol transport machinery, to reach the extracellular space bypassing the Golgi complex. A similar pathway appears to be true for the ZIKV NS1. These results are in congruence with the lipoprotein nature of NS1 and uncover novel functions for the CCC as a protein transporter in mosquito cells. The presence of the CBD in NS1 appears to be important to mediate the interaction between NS1 and CAV-1. Yet, more research in needed to fully understand the viral and cellular factors that determinate the dramatic differences in traffic route followed by DENV NS1 in mosquito and in vertebrate cells. NS1 proteins from others mosquito borne flaviviruses that lack the CBD, as WNV, YFV, JEV are secreted apparently at very low levels and following the classical pathway in mosquito cells. The functions of soluble NS1 in the mosquito are unknown and need to be elucidated; but may include facilitation of the spread of viral particles and modulation of innate immunity [31]. Finally, manipulation of the lipid and cholesterol system transport in the mosquito cell may become a novel target to reduce NS1 secretion and a novel strategy to blocks the spread of mosquito-borne flaviviruses such as dengue virus.

## MATERIALS AND METHODS

### Cells

C6/36 clone mosquito cells from *Aedes albopictus* (ATCC^®^ CRL-1660™) and Aag-2 cells from *Aedes aegypti* (kindly provided by Dr. Fidel de la Cruz from CINVESTAV-IPN) were grow at 28 °C in Eagle’s Minimum Essential Medium (EMEM) (ATCC^®^ 30-2003™), supplemented with 5% fetal bovine serum (FBS) and 100 U/ml penicillin-streptomycin. Baby hamster kidney cells (BHK-21, ATCC^®^ CCL-10™) were grown at 37°C and cultured in Eagle’s Minimum Essential Medium (EMEM, ATCC^®^) supplemented with 5% FBS and 100 U/ml penicillin-streptomycin. Monkey epithelial kidney cells Vero E6 (ATCC^®^ CRL-1586™) were grown at 37°C and cultured in Eagle’s Minimum Essential Medium (EMEM, ATCC^®^) supplemented with 10% FBS and 100 U/ml penicillin-streptomycin.

### Viruses

DENV serotype 2 (DENV2), DENV serotype 4 (DENV4) and Zika virus (ZIKV), were generously provided by Msc. Mauricio Vázquez, (Laboratorio de Arbovirus y Virus Hemorrágicos. Instituto de Diagnóstico y Referencia Epidemiológicos (InDRE, Mexico City). The dengue viruses were propagated in suckling mice brain (ICR(CD-1), provided by Unit of production and experimentation of laboratory animals of CINVESTAV-IPN, Mexico (UPEAL-CINVESTAV-IPN) as previously described [52]. Virus titers in mice brain homogenates were determined by focus forming assay in BHK-21 cells. Serial dilutions of viral stocks or supernatants of experiments were diluted in serum-free medium and added to cell monolayers in 96-well plates. Viral attachment and entry were allowed to proceed for 2 h at 37°C; then EMEM 10% FBS was added and incubated for an additional 48 h before fixation and quantitation of infected cells by labelling with Anti-Flavivirus E-glycoprotein antibody using mouse VECTASTAIN^®^ ABC-HRP Kit (PK-4002-VectorLabs) and DAB Peroxidase Substrate Kit (SK-4100-Vector Labs). ZIKV was propagated in Vero E6 cells. YFV-17DD vaccine strain stock was donated by Dr. Juan Salas Benito (Escuela Nacional de Medicina y Homeopatía, IPN) and obtained from passages of the commercial vaccine in C6/36 cells. Infection experiments with YFV and ZIKV and stock viruses were tittered in Vero E6 cells using focus forming assay (as described previously).

### Luciferase reporter construct

Ac5-STABLE2-neo was a gift from Rosa Barrio and James Sutherland (Addgene plasmid # 32426) [53]. This plasmid was engineered to express Luciferase gene in mosquito cells. Luciferase gene was obtained from plasmid ptk-GLuc Vector (NEB, Catalog # N8084) kindly donated by Dr. Susana Lopez (Instituto de Biotecnologia-UNAM). *Gaussia* Luciferase (GLuc) was PCR-amplified using the following designed primers with directed cloning sites (underlined): Luc-XbaI-F (5’-GCGC<u>TCTAGA</u>GCCACCATGGGAGTCAAAGTTCTGTTTGCC-3’) and Luc-HindIII-R primer (5’-GAGT<u>AAGCTTT</u>GGCGGGTCACCACCGGCCCCCTTGATC-3’). Unique sites XbaI and HindIII in pAc5-stable2-neo allowed replacement of GFP. XbaI-HindIII cassette was cloned into pAc5-STABLE2-Neo, generating the pAc5–mCherry-GLuc-Neo vector. The promoter Actin5C (from *Drosophila melanogaster*) is efficient in insect cells lines. The T2A peptide sequence derived from *Thosea asigna* (EGRGSLLTCGDVEENPGP) allowed multicistronic processing and the neomycin resistance gene (NeoR) confers resistance to G418 allowing stable mosquito cell lines. Schematic representation of pAc5-mCherry-GLuc-neo vector is shown in S1 Fig.

### Vector transfection and Luciferase activity assay

pAc5-mCherry-GLuc-neo vector was transfected into confluent monolayer of C6/36 cells using lipofectamine reagent LipofectamineTM2000 (Invitrogen). Each 24-well was transfected with 1 μg of plasmid DNA and 2 μL of Lipofectamine. After 5 h of transfection, cells were added EMEM with a final 10% FBS. After 24h, selective G418 was added to obtain stable C6/36 cells line. Stable C6/36 cells were treated with BFA or GCA for 24h. BHK-21 cells were transfected with pkt-GLuc vector and drug treatment was following the same procedure detailed above. Subsequently, the supernatants were analyzed with BioLux^®^ *Gaussia* Luciferase Assay Kit (NEB). Percentage of luciferase activity was normalized with DMSO treatment.

### Reagents and drug treatment

Brefeldin A (BFA) (B6542-Sigma-Aldrich), Golgicide A (GCA, 1139889-93-2 Calbiochem), Fli-06 (SML0975, Sigma-Aldrich) and Cyclosporine A (CsA, C1832-Sigma-Aldrich) were dissolved in dimethyl sulfoxide (DMSO, ATCC^®^). CsA was used to pharmacologically inhibit Cyclophilin A. Methyl-β-cyclodextrin (MβCD, C4555-Sigma-Aldrich) was dissolved in water and used to inhibit mobilization of cholesterol and to alters lipids within cells. Unless indicated otherwise, the experimental concentrations of BFA, GCA, Fli-06, CsA and MβCD were 7 μM, 27 μM, 100 μM, 9 μM and 9 mM, respectively. Cells grown in 24-well plates were infected for 1 hour and then washed three times with PBS. The drugs BFA, GCA, CsA and Fli-06 were added to the cells in EMEM 5% FBS. Unless indicated otherwise, incubation time was 24 hours at 28°C for mosquito cells and 37°C for BHK-21 cells. After this time, cell supernatants were collected to measure secreted NS1 and virus progeny. In other cases, cells were fixed and stained for immunofluorescence.

### Cell viability assays

After treatment with the indicated drugs or siRNAs, the number of viable cells in each sample was determined in triplicate using the Cell Titer 96 AQueous Non-Radioactive Cell Proliferation Assay according to the manufacturing procedures (MTS assay, G3580-Promega). Triplicate measurements were then averaged and the percentage of viable cells determined relative to cells treated as controls.

### Golgi integrity after drug treatment

Cells grown to subconfluency on coverslips in 24-well plates were treated with GCA or Fli-06 for 24h. After fixation with 4% paraformaldehyde, the cells were stained with rabbit monoclonal anti-GM130 antibody (Sigma-Aldrich). The samples were stained with the secondary antibody Alexa Fluor 488-conjugated and Golgi apparatus integrity was analyzed with a Nikon Epifluorecence microscope (Eclipse Ti-U).

### Measurement of secreted NS1 protein

The PlateliaTM NS1 Ag (Bio-Rad, #72830 Hercules, CA) commercial kit was used to determine the levels of soluble NS1 in cell supernatants collected from silenced, drug treated or DENV infected cells. The assay was carried out following the procedure indicated by the manufacturer. The secreted NS1 was reported as the percentage of untreated or mock infected cells.

### Gene knockdown with siRNAs

C6/36 and BHK-21 cells were transfected with a range of concentrations of siRNA targeted against proteins of the chaperon caveolin complex using HiPerFect Transfection Reagent (QIAGEN^®^). FlexiTube siRNA (4 siRNAs recommended per each gene, QIAGEN^®^) for FKBP52, CAV-1 and Cy40 were used to concentrations indicated in text. AllStars Negative Control siRNA (QIAGEN^®^) was used as non-silencing siRNA control at the same concentration of the siRNAs for the proteins of interest. Gene knock down was assessed using Western Blotting. Protein expression was normalized to the expression of β-actin (CINVESTAV-IPN), GAPDH (GTX100118, Genetex^®^) or ERK-1 (sc-271291, Santa Cruz Biotechnology, Inc). Expression levels relative to negative control cells were estimated by densitometric measuring with ImageJ software [54].

### Lipid droplet count

Lipid droplet count per cell was used as an indicator to determine the effect on lipid homeostasis by drug or siRNA treatment. Cells were fixed and washed three times in PBS. Stock solution of Nile Red (72485 Sigma-Aldrich) 500 μg/ml in acetone was prepared and stored protected from light. The dye was then added directly to the preparation at 0.5 μg/ml in PBS, and the preparation was incubated for 30 min. After three washes in PBS excess of dye was removed. From here in advance all suspension medium did not contain serum, albumin or detergents to avoid draining the dye out of the cells. Then, coverslips were mounted in Fluoroshield™ with DAPI (Sigma). Lipid droplets were numbered in maximum projection images obtained by LSM 700 confocal microscope. The total amount of lipid droplets was determined in at least 15 cells using Spot Detector plugin [55] at Icy Image software (Institute Pasteur) [56].

### Confocal microscopy

Confluent cell monolayers, grown in 24-well plates containing glass coverslips, were infected with DENV infected using a MOI=3. After the times indicated in the text, cells were fixed in paraformaldehyde 4% for 10 min. Cells were permeabilizated with 0.1% Triton X-100 for 10 minutes at room temperature and stained for DENV-NS1 using anti-NS1 2B7, anti-GM130 (G7295 Sigma-Aldrich), anti-CAV-1 (GTX89541 Genetex or sc-894 Santa Cruz), anti-FKBP52 (ab129097 Abcam), anti-Cy40 (GTX104038 Genetex), anti-CyA (GTX104698 Genetex) and Nuclei with DAPI. Anti-mouse Alexa-488 or Alexa-598, anti-goat Alexa-568 and Anti-rabbit Alexa-647 or Alexa-488 conjugated (Donkey pre-adsorbed, secondary antibodies, Abcam) were used at 1:800 dilution). For lipid droplets co-localization with DENV NS1, cells were incubated with BODIPY 493/503 (Invitrogen) at 2 μM, in PBS-free of serum or detergents. Coverslips were mounted in Fluoroshield™ with DAPI (Sigma). The images were analyzed using a Carl Zeiss LSM 700 confocal microscope. To maintain the consistency of the green color for the viral protein, the color of BODIPY was changed to red as for Nile red staining. To evaluate the co-localization between proteins, Pearson correlation coefficients (PCC) were obtained from at least 20 confocal independent images (laser sections indicated in text) using the Icy image software and the co-localization studio plugin.

### Western blotting

Total cells were lysed in Lysis Buffer (25mM Tris-HCl pH 7.6, 150mM NaCl, 1% Triton X-100, 1% sodium deoxycholate, 0.1% SDS, 5% Glycerol) with protease inhibitor cocktail (P8340 Sigma) added and assayed for total protein concentration using Pierce™ BCA Protein Assay Kit (Thermo Scientific). Samples were subsequently prepared for electrophoresis by adding 4X Laemmli gel loading buffer (40% Glycerol, 240 mM Tris-HCl pH 6.8, 8% SDS, 0.04% bromophenol blue, 5% β-mercaptoethanol). Samples were then boiled and resolved in SDS-polyacrylamide gels, transferred to nitrocellulose membrane 0.45 μm (Bio-Rad), and incubated with primary antibodies diluted in 5% skin milk powder in PBS-0.1% Tween-20. After washing, membranes were incubated with secondary antibodies conjugated to horseradish peroxidase (HRP) (anti-mouse-HRP 115-035-003 Jackson ImmunoResearch or anti-rabbit-HRP GTX26721 Genetex) diluted in 5% skin milk powder in PBS −0.1% Tween-20. HRP was detected using SuperSignal™ West Femto Maximum Sensitivity Substrate (Thermo Scientific). Digital images was acquired with a Fusion FX Spectra (Vilber) and analyzed with ImageJ software [54].

### Immunoprecipitation

Lysates was harvested from DENV infected cells using ice-cold non-denaturing lysis buffer IP (PBS pH 7.4, 1% Triton X-100, NaCl 150 mM, Glycerol 5% and protease inhibitor cocktail). Co-immunoprecipitation (co-IP) was done using the Thermo Scientific Pierce co-IP kit following the manufacturer’s protocol. Briefly, the DENV NS1 polyclonal antibody (GTX124280 Genetex) and anti-CAV-1 polyclonal (n-20 Santa cruz biotechnology) were first immobilized for 2 h using AminoLink Plus coupling resin. The resin was then washed and incubated with the cell lysates overnight. After incubation, the resin was again washed (PBS pH 7.4, NaCl 150 mM, glycerol 5%) and proteins eluted using elution buffer (Hepes 50mM pH 5.0). A mock infection control was used to assess nonspecific binding, received the same treatment as the co-IP samples, including the CAV-1 antibody. In this control, the coupling resin is not amine-reactive preventing covalent immobilization of the primary antibody onto the resin. Samples were analyzed by immunoblotting using anti-CAV-1, anti-FKBP52, anti-Cy40, anti-CyA and anti-dengue NS1 (antibodies previously described). Membranes were then incubated with HRP-conjugated secondary antibodies (anti-rabbit-HRP or anti-mouse-HRP) and developed using enhanced chemiluminescence (Thermo Scientific).

### Proximity ligation assay

Interactions between the NS1 and the CCC proteins were detected by Duolink proximity ligation assay (PLA) Kit (Sigma Aldrich) in DENV4-infected mosquito cells (MOI=3). After 18hpi infection, cells were fixed and permeabilizated in methanol. After pre-incubation with a blocking agent for 1 h, samples were incubated overnight with the primary antibodies overnight. Duolink PLA probes detecting rabbit or mouse antibodies were diluted in the blocking agent a dilution of 1:5 and applied to the slides, followed by incubation for 1 h in a pre-heated humidity chamber at 37°C. The PLA probe anti-rabbit Plus binds to the CAV-1, Cy40, CyA or FKBP52 primary antibodies, whereas the PLA probe anti-mouse Minus binds to the dengue NS1 antibody. If the distance between both probes is <40 nm, a signal with DuoLink PLA is generated and detected in FITC dye emission, thus indicating an interaction of both proteins. Unbound PLA probes were removed by washing in buffer A (10 mM Tris-HCl pH 7.4, 150mM NaCl, 0.05% Tween 20). For hybridization of the two Duolink PLA probes, Duolink Hybridization stock (dilution 1:5) was used. Slides were incubated in a pre-heated humidity chamber for 15 min at 37°C. The samples were incubated in the ligation solution consisting of Duolink Ligation stock (1:5) and Duolink Ligase (1:40) for 90 min at 37°C. Detection of the amplified probe was done with the Duolink Detection Green Kit. Duolink Detection stock was diluted at 1:5 and applied for 1 h at 37°C. Final washing steps were done in buffer B (200mM Tris-HCl pH 7.5, and 100mM NaCl). Then, the slides were mounted with *In Situ* Mounting Medium with DAPI (Sigma) and visualized by LSM 700 confocal microscope. The PLA signals per cell were determined in at least 10 cells in maximum projection images using Spot Detector plugin at Icy Image software [56]. Mock infected cells incubated with both primary antibodies or DENV infected cells incubated only with anti-CAV-1 antibodies were included as negative controls.

### Protein-protein interaction *in silico* analysis

3D crystal structures of dimeric dengue NS1 (PDB: 4OIG) and proteins of the CCC, FKBP52 (4LAV), CyA (1OCA) and Cy40 (1IHG), were taken from PDB databank and analyzed using ClusPro protein-protein docking experiments server (https://cluspro.bu.edu/home.php). The highest favorable binding energies were obtained by models using scoring schemes in hydrophobic interactions. Sixteen clusters of low-energy docked structures were analyzed with each energy parameter and compared with dimeric dengue NS1. Since no amino acid sequence from CAV-1 of any mosquito species is annotated and no crystal structure for the human CAV-1 is found in the PDB data bank, the CAV-1 putative 3D structure based on the amino acid sequence of human CAV-1 (isoform alpha, NP_001744.2) was modeled using a protein structure prediction server [57]. Then, CAV-1 putative structure was used in ClusPro protein-protein docking analysis with NS1 and proteins of CCC [58]. Dockings were performed with the balanced free energy function and the values were statistically analyzed and compared [25]. 3D structures and atom to atom distances were represented and measured using PyMOL software (The PyMOL Molecular Graphics System, Version 2.0 Schrödinger, LLC).

### Statistical analysis

Values of all assays were expressed as mean ± standard error of three independent experiments, each in triplicate or indicated in the text. Statistical analyzes were carried out using the GraphPad Prism version 6.01 software.

## ACKNOWLEDGMENTS

Authors like to thank Ana C. Alcalá for helpful discussions about results and experiments.

## SUPPLEMENTAL FIGURES

**S1 Fig. Schematic representation of the pAc5-mCherry-GLuc-neo vector.**

pAc5-mCherry-GLuc-Neo was constructed by cloning GLuc into the XbaI-HindIII sites of pAc5-STABLE2-Neo designed for tricistronic expression driven by the Actin5C promoter. Cloning sites are shown. Features: FLAG-tagged mCherry, *Gaussia* Luciferase and NeoR, each separated by a T2A peptide.

**S2 Fig. DENV NS1 does not co-localizes of with lipid droplets**

Co-localization of NS1 and lipid droplets in mosquito DENV2 and DENV4 infected cells. LD were stained with Bodipy and shown in red, DENV NS1 is shown in green and nuclei in blue (DAPI). The lateral small panels show magnification of the selected zones. Pearson correlation coefficient (PCC) was used to evaluate the degree of co-localization between NS1 and LD. PCC are shown on top of each square panels. The images were analyzed using a LSM 700 confocal microscope with laser sections: 0.45 μm.

**S3 Fig. Lipid droplet counting in C6/36 cells after cholesterol depletion**

DENV4 infection (MOI=3), MβCD treatment (9 mM), and starvation was performed in absence of serum. Lipid droplets (LD) were stained with Nile red 24h after each treatment. LD were counted in maximum projection images in treated cells from at least 20 cells by group, using spot detector plugin using Icy software. LD median count differences were compared with Mock using One way-ANOVA denoted by * (P ≤ 0.05).

**S4 Fig. Chaperone caveolin complex co-localization in non-infected vertebrate and mosquito cells.**

**A**, **B,** and **C** Co-localization for NS1 and CCC in C6/36 cells. **D**, **E,** and **F** Co-localization for NS1 and CCC in Aag2 cells. ***G***, ***H,*** *and* ***I*** Co-localization for NS1 and CCC in BHK-21cells. Mock cells were fixed and probed against CAV-1 (shown in red) and FKBP52, CyA, and Cy40 (shown in green). Merged fluorescent images of DAPI, red and green channels are shown. The images were analyzed using a LSM 700 confocal microscope. Pearson correlation coefficients (PCC) were used to analyze degree of co-localization with laser sections: 0.41 μm (PCC were shown in Fig 7N).

**S5 Fig. ZIKV and YFV virion release decrease in cells treated with classical secretion pathway inhibitors**

**A** Viral release in YFV infected cells treated with inhibitors of classical secretion pathway at 72 hpi. **B** Viral release in ZIKV infected cells treated with inhibitors of classical secretion pathway at 48 hpi. ***A*** *and* ***B*** BFA and GCA treatment at 7 and 27 μM, respectively. Viral titers in supernatants were measured by focus forming assay in Vero E6 cells according to standard procedures. The experiments were performed in triplicate, and the bars represent means ± standard error. Data was evaluated using the 2way ANOVA test and significant differences compared with DMSO treatment group, respectively, are denoted by *(P ≤0.05).

## REFERENCES

1. Guzman MG, Harris E. Dengue. Lancet. 2015;385: 453–465. doi:10.1016/S0140-6736(14)60572-9Seminar

2. Bhatt S, Gething PW, Brady OJ, Messina JP, Farlow AW, Moyes CL, et al. The global distribution and burden of dengue. Nature. 2013;496: 504–507. doi:10.1038/nature12060

3. Acosta EG, Kumar A, Bartenschlager R. Revisiting Dengue Virus–Host Cell Interaction. Advances in virus research. 2014. pp. 1–109. doi:10.1016/B978-0-12-800098-4.00001-5

4. Muller DA, Young PR. The flavivirus NS1 protein: Molecular and structural biology, immunology, role in pathogenesis and application as a diagnostic biomarker. Antiviral Res. Elsevier; 2013;98: 192–208. doi:10.1016/J.ANTIVIRAL.2013.03.008

5. Akey ADL, Brown WC, Dutta S, Konwerski J, Jose J. Flavivirus NS1 crystal structures reveal a surface for membrane association and regions of interaction with the immune system. Science (80-). 2014;343: 1–31. doi:10.1126/science.1247749.Flavivirus

6. Gutsche I, Coulibaly F, Voss JE, Salmon J, d’Alayer J, Ermonval M, et al. Secreted dengue virus nonstructural protein NS1 is an atypical barrel-shaped high-density lipoprotein. Proc Natl Acad Sci. 2011;108: 8003–8008. doi:10.1073/pnas.1017338108

7. Alcalá AC, Medina F, González-Robles A, Salazar-Villatoro L, Fragoso-Soriano RJ, Vásquez C, et al. The dengue virus non-structural protein 1 (NS1) is secreted efficiently from infected mosquito cells. Virology. Elsevier; 2016;488: 278–287. doi:10.1016/j.virol.2015.11.020

8. Thiemmeca S, Tamdet C, Punyadee N, Prommool T, Songjaeng A, Noisakran S, et al. Secreted NS1 Protects Dengue Virus from Mannose-Binding Lectin–Mediated Neutralization. J Immunol. 2016;197: 4053–4065. doi:10.4049/jimmunol.1600323

9. Flamand M, Megret F, Mathieu M, Lepault J, Rey FA, Deubel V. Dengue virus type 1 nonstructural glycoprotein NS1 is secreted from mammalian cells as a soluble hexamer in a glycosylation-dependent fashion. J Virol. American Society for Microbiology; 1999;73: 6104–10. Available: http://www.ncbi.nlm.nih.gov/pubmed/10364366

10. Alcalá AC, Hernández-Bravo R, Medina F, Coll DS, Zambrano JL, Del Angel RM, et al. The dengue virus non-structural protein 1 (NS1) is secreted from infected mosquito cells via a non-classical caveolin-1-dependent pathway. J Gen Virol. 2017;98: 2088–2099. doi:10.1099/jgv.0.000881

11. Parton RG, Simons K. The multiple faces of caveolae. Nat Rev Mol Cell Biol. 2007;8: 185–194. doi:10.1038/nrm2122

12. Fielding CJ, Fielding PE. Cholesterol and caveolae: Structural and functional relationships. Biochim Biophys Acta - Mol Cell Biol Lipids. 2000;1529: 210–222. doi:10.1016/S1388-1981(00)00150-5

13. Rothberg KG, Heuser JE, Donzell WC, Ying Y-S, Glenney JR, Anderson RGW. Caveolin, a protein component of caveolae membrane coats. Cell. Cell Press; 1992;68: 673–682. doi:10.1016/0092-8674(92)90143-Z

14. Razani B, Woodman SE, Lisanti MP. Caveolae: from cell biology to animal physiology. Pharmacol Rev. 2002;54: 431–467. doi:10.1124/pr.54.3.431

15. Dawar FU, Xiong Y, Khattak MNK, Li J, Lin L, Mei J. Potential role of cyclophilin A in Regulating cytokine secretion. J Leukoc Biol. 2017;102: 989–992. doi:10.1189/jlb.3RU0317-090RR

16. Wang P, Heitman J. The cyclophilins. Genome Biol. 2005;6: 226. doi:10.1186/gb-2005-6-7-226

17. Guy NC, Garcia YA, Cox MB. Therapeutic Targeting of the FKBP52 Co-Chaperone in Steroid Hormone Receptor-Regulated Physiology and Disease. Curr Mol Pharmacol. 2015; 109–125.

18. Li P, Ding Y, Wu B, Shu C, Shen B, Rao Z. Structure of the N-terminal domain of human FKBP52. Acta Crystallogr D Biol Crystallogr. 2003;59: 16–22. Available: http://www.ncbi.nlm.nih.gov/pubmed/12499534

19. Waldmeier PC, Zimmermann K, Qian T, Tintelnot-Blomley M, Lemasters JJ. Cyclophilin D as a drug target. Curr Med Chem. 2003;10: 1485–506. Available: http://www.ncbi.nlm.nih.gov/pubmed/12871122

20. van der Linden L, van der Schaar HM, Lanke KHW, Neyts J, van Kuppeveld FJM. Differential Effects of the Putative GBF1 Inhibitors Golgicide A and AG1478 on Enterovirus Replication. J Virol. 2010;84: 7535–7542. doi:10.1128/JVI.02684-09

21. Krämer A, Mentrup T, Kleizen B, Rivera-Milla E, Reichenbach D, Enzensperger C, et al. Small molecules intercept Notch signaling and the early secretory pathway. Nat Chem Biol. 2013;9: 731–738. doi:10.1038/nchembio.1356

22. Yonemura Y, Li X, Müller K, Krämer A, Atigbire P, Mentrup T, et al. Inhibition of cargo export at ER exit sites and the trans-Golgi network by the secretion inhibitor FLI-06. J Cell Sci. 2016;129: 3868–3877. doi:10.1242/jcs.186163

23. Guy NC, Garcia YA, Sivils JC, Galigniana MD, Cox MB. Functions of the Hsp90-binding FKBP immunophilins [Internet]. Sub-cellular biochemistry. 2015. pp. 35–68. doi:10.1007/978-3-319-11731-7_2

24. Ricciardi-Jorge T, Bordignon J, Koishi A, Zanluca C, Mosimann AL, Duarte dos Santos CN. Development of a quantitative NS1-capture enzyme-linked immunosorbent assay for early detection of yellow fever virus infection. Sci Rep. 2017;7: 16229. doi:10.1038/s41598-017-16231-6

25. Wang Z, Xu J. Predicting protein contact map using evolutionary and physical constraints by integer programming (extended version). 2013; doi:10.1093/bioinformatics/btt211

26. Stepanyuk GA, Xu H, Wu C-K, Markova S V, Lee J, Vysotski ES, et al. Expression, purification and characterization of the secreted luciferase of the copepod Metridia longa from Sf9 insect cells. Protein Expr Purif. NIH Public Access; 2008;61: 142–8. doi:10.1016/j.pep.2008.05.013

27. Henderson J, Macdonald H, Lazarus CM, Napier RM, Hawes CR. Protein retention in the endoplasmic reticulum of insect cells is not compromised by baculovirus infection. Cell Biol Int. Wiley-Blackwell; 1996;20: 413–422. doi:10.1006/cbir.1996.0052

28. Youn S, Cho H, Fremont DH, Diamond MS. A Short N-Terminal Peptide Motif on Flavivirus Nonstructural Protein NS1 Modulates Cellular Targeting and Immune Recognition. J Virol. 2010;84: 9516–9532. doi:10.1128/JVI.00775-10

29. Imai S, Kusakabe T, Xu J, Li Z, Shirai S, Mon H, et al. Roles of silkworm endoplasmic reticulum chaperones in the secretion of recombinant proteins expressed by baculovirus system. Mol Cell Biochem. 2015;409: 255–262. doi:10.1007/s11010-015-2529-5

30. Teng C-Y, Chang S-L, van Oers MM, Wu T-Y. Enhanced Protein Secretion From Insect Cells by Co-Expression of the Chaperone Calreticulin and Translation Initiation Factor eIF4E. Mol Biotechnol. 2013;54: 68–78. doi:10.1007/s12033-012-9545-4

31. Gullberg M, Andersson A-C. Visualization and quantification of protein-protein interactions in cells and tissues. Nat Methods. Nature Publishing Group; 2010;7: v–vi. doi:10.1038/nmeth.f.306

32. Henkhaus RS, Roy UKB, Cavallo-Medved D, Sloane BF, Gerner EW, Ignatenko NA. Caveolin-1-Mediated Expression and Secretion of Kallikrein 6 in Colon Cancer Cells. Neoplasia. Elsevier; 2008;10: 140–148. doi:10.1593/NEO.07817

33. Jin Y, Kim HP, Cao J, Zhang M, Ifedigbo E, Choi AMK. Caveolin-1 regulates the secretion and cytoprotection of Cyr61 in hyperoxic cell death. FASEB J. Federation of American Societies for Experimental Biology; 2009;23: 341–350. doi:10.1096/fj.08-108423

34. Mir KD, Parr RD, Schroeder F, Ball JM. Rotavirus NSP4 interacts with both the amino-and carboxyl-termini of caveolin-1. Virus Res. NIH Public Access; 2007;126: 106–115. doi:10.1016/j.virusres.2007.02.004

35. Ball JM, Schroeder ME, Williams C V, Schroeder F, Parr RD. Mutational analysis of the rotavirus NSP4 enterotoxic domain that binds to caveolin-1. Virol J. 2013;10: 336. doi:10.1186/1743-422X-10-336

36. Khachatoorian R, French SW. Chaperones in hepatitis C virus infection. World J Hepatol. 2016;8: 9–35. doi:10.4254/wjh.v8.i1.9

37. Gao J, Xiao S, Liu X, Wang L, Zhang X, Ji Q, et al. Inhibition of HSP90 attenuates porcine reproductive and respiratory syndrome virus production in vitro. Virol J. Virology Journal; 2014;11: 17. doi:10.1186/1743-422X-11-17

38. Das I, Basantray I, Mamidi P, Nayak TK, Pratheek BM, Chattopadhyay S, et al. Heat shock protein 90 positively regulates Chikungunya virus replication by stabilizing viral non-structural protein nsP2 during infection. PLoS One. 2014;9. doi:10.1371/journal.pone.0100531

39. Srisutthisamphan K, Jirakanwisal K, Ramphan S, Tongluan N, Kuadkitkan A, Smith DR. Hsp90 interacts with multiple dengue virus 2 proteins. Sci Rep. Nature Publishing Group; 2018;8: 4308. doi:10.1038/s41598-018-22639-5

40. Dawar FU, Tu J, Khattak MNK, Mei J, Lin L. Cyclophilin a: A key factor in virus replication and potential target for anti-viral therapy. Curr Issues Mol Biol. 2017;21: 1–20.

41. Yang F, Robotham JM, Nelson HB, Irsigler A, Kenworthy R, Tang H. Cyclophilin A is an essential cofactor for hepatitis C virus infection and the principal mediator of cyclosporine resistance in vitro. J Virol. 2008;82: 5269–78. doi:10.1128/JVI.02614-07

42. Tanaka Y, Sato Y, Sasaki T. Suppression of coronavirus replication by cyclophilin inhibitors. Viruses. 2013;5: 1250–1260. doi:10.3390/v5051250

43. Favreau DJ, Meessen-Pinard M, Desforges M, Talbot PJ. Human Coronavirus-Induced Neuronal Programmed Cell Death Is Cyclophilin D Dependent and Potentially Caspase Dispensable. J Virol. 2012;86: 81–93. doi:10.1128/JVI.06062-11

44. Quarato G, D’Aprile A, Gavillet B, Vuagniaux G, Moradpour D, Capitanio N, et al. The cyclophilin inhibitor alisporivir prevents hepatitis C virus-mediated mitochondrial dysfunction. Hepatology. 2012;55: 1333–1343. doi:10.1002/hep.25514

45. Perera R, Riley C, Isaac G, Hopf-Jannasch AS, Moore RJ, Weitz KW, et al. Dengue virus infection perturbs lipid homeostasis in infected mosquito cells. PLoS Pathog. Public Library of Science; 2012;8: e1002584. doi:10.1371/journal.ppat.1002584

46. Rolls MM, Marquardt MT, Kielian M, Machamer CE. Cholesterol-independent targeting of Golgi membrane proteins in insect cells. Mol Biol Cell. American Society for Cell Biology; 1997;8: 2111–8. Available: http://www.ncbi.nlm.nih.gov/pubmed/9362056

47. Vinci G, Xia X, Veitia RA. Preservation of Genes Involved in Sterol Metabolism in Cholesterol Auxotrophs: Facts and Hypotheses. Volff J-N, editor. PLoS One. Public Library of Science; 2008;3: e2883. doi:10.1371/journal.pone.0002883

48. Soto-Acosta R, Bautista-Carbajal P, Cervantes-Salazar M, Angel-Ambrocio AH, del Angel RM. DENV up-regulates the HMG-CoA reductase activity through the impairment of AMPK phosphorylation: A potential antiviral target. Kuhn RJ, editor. PLOS Pathog. 2017;13: e1006257. doi:10.1371/journal.ppat.1006257

49. Samsa MM, Mondotte JA, Iglesias NG, Assunção-Miranda I, Barbosa-Lima G, Da Poian AT, et al. Dengue virus capsid protein usurps lipid droplets for viral particle formation. PLoS Pathog. 2009;5. doi:10.1371/journal.ppat.1000632

50. Leier HC, Messer WB, Tafesse FG. Lipids and pathogenic flaviviruses: An intimate union. Dutch RE, editor. PLOS Pathog. Public Library of Science; 2018;14: e1006952. doi:10.1371/journal.ppat.1006952

51. Byrne DP, Dart C, Rigden DJ. Evaluating Caveolin Interactions: Do Proteins Interact with the Caveolin Scaffolding Domain through a Widespread Aromatic Residue-Rich Motif? PLoS One. 2012;7. doi:10.1371/journal.pone.0044879

52. Gould EA CJ. Growth, titulation and purification of alphaviruses and flaviviruses. Virology: a practical approach. 1991. pp. 43–78.

53. González M, Martín-Ruíz I, Jiménez S, Pirone L, Barrio R, Sutherland JD. Generation of stable Drosophila cell lines using multicistronic vectors. Sci Rep. 2011;1. doi:10.1038/srep00075

54. Rueden CT, Schindelin J, Hiner MC, DeZonia BE, Walter AE, Arena ET, et al. ImageJ2: ImageJ for the next generation of scientific image data. BMC Bioinformatics. BioMed Central; 2017;18: 529. doi:10.1186/s12859-017-1934-z

55. Olivo-Marin J-C. Extraction of spots in biological images using multiscale products. Pattern Recognit. Pergamon; 2002;35: 1989–1996. doi:10.1016/S0031-3203(01)00127-3

56. de Chaumont F, Dallongeville S, Chenouard N, Hervé N, Pop S, Provoost T, et al. Icy: an open bioimage informatics platform for extended reproducible research. Nat Methods. 2012;9: 690–696. doi:10.1038/nmeth.2075

57. Ma J, Wang S, Zhao F, Xu J. Protein threading using context-specific alignment potential. Bioinformatics. Oxford University Press; 2013;29: i257–i265. doi:10.1093/bioinformatics/btt210

58. Kozakov D, Hall DR, Xia B, Porter KA, Padhorny D, Yueh C, et al. The ClusPro web server for protein-protein docking. Nat Protoc. 2017;12: 255–278. doi:10.1038/nprot.2016.169

